# Effect heterogeneity reveals complex pleiotropic effects of rare coding variants

**DOI:** 10.1101/2024.10.01.614806

**Authors:** Wenhan Lu, Siwei Chen, Chiara Auwerx, Jack Fu, Danielle Posthuma, Benjamin M. Neale, Luke J. O’Connor, Konrad J. Karczewski

## Abstract

Recent expansion of large-scale biobank resources has enabled rare-variant association studies (RVAS) and systematic investigation of rare variant pleiotropy across thousands of phenotypes simultaneously. However, existing statistical frameworks for dissecting pleiotropy were largely developed for common variants and are not well suited to gene-based rare variant signals, limiting the interpretation of cross-phenotype associations. Here, we develop ALLSPICE, a likelihood-based method that tests whether rare variant effects in genes exhibiting cross-phenotype associations are proportional or heterogeneous across continuous traits while accounting for phenotypic correlation using summary statistics. ALLSPICE is well-calibrated in simulations and is implemented as an R package for scalable analysis of gene-level rare-variant burden associations. We applied ALLSPICE to RVAS of 359 continuous traits in the UK Biobank and identified 124 significant heterogeneous events among 11,810 pairs of gene-trait associations. By identifying effect heterogeneity within genes associated with multiple phenotypes, ALLSPICE clarifies shared and heterogeneous rare variant architectures underlying cross-phenotype associations and provides insight into rare variant pleiotropy.

## Introduction

Genome-wide association studies (GWAS) have identified thousands of loci associated with multiple phenotypes, indicating that shared genetic effects are widespread across complex traits^1–3^. Such cross-phenotype associations may arise through different forms of pleiotropy, in which a single gene, variant, or genetic perturbation affects multiple phenotypes^4^. For example, vertical pleiotropy occurs when the genetic effect on one trait causally mediates its effect on another trait and is commonly evaluated using causal inference approaches such as Mendelian Randomization (MR)^5^. By contrast, horizontal pleiotropy refers to independent effects of a shared genetic component on multiple traits through distinct biological mechanisms^6^, as illustrated by variation in *MC1R* influencing multiple pigmentation phenotypes^7^.

Although pleiotropy has been extensively studied for common variants^6,8–13^, similar inference for rare variants remains largely unexplored. Existing methods for dissecting pleiotropy were primarily developed for common variants and rely on assumptions that do not readily extend to rare variants, where limited single-variant power generally requires gene-level aggregation. In rare variant association studies (RVAS), gene-level burden signals arise from sparse allelic architectures in which rare variants are grouped by frequency and predicted functional class, but residual heterogeneity in variant-level effects can still strongly influence the resulting association signal^14^. While pleiotropic rare variant associations have been reported within specific phenotype groups^15–18^, the contribution of heterogeneous allelic effects to gene-burden signals has not been systematically characterized across the phenome.

To address this gap, we developed ALLSPICE (ALLelic Spectrum of Pleiotropy Informed Correlated Effects), a likelihood-based framework for testing whether rare variant effects in genes with cross-phenotype burden associations are consistent with proportional or heterogeneous variant-level architectures across continuous traits. Under a shared mediation model, variant effects are expected to remain approximately proportional across phenotypes, reflecting a common causal pathway, whereas deviations from proportionality suggest divergent contributions from different allelic subsets. Specifically, for a gene exhibiting burden associations with two traits, ALLSPICE tests whether observed variant-level effects within the gene are compatible with a proportional model, *H*_0_ : β_1_ = *c*β_2_ across the two traits, where *c* is estimated by maximum likelihood under a closed-form linear regression framework that accounts for phenotypic correlation. Because this likelihood is derived under a Gaussian linear model, the current implementation is restricted to continuous traits. ALLSPICE is designed for ultra-rare variants (minor allele frequency < 0.01%), for which linkage disequilibrium is expected to be limited, and variant effects can be treated as approximately independent at the summary-statistic level.

ALLSPICE is computationally efficient and implemented in the R package *ALLSPICER*. The method uses single-variant association summary statistics and therefore can be applied directly to large RVAS resources without requiring individual-level genotype data. In simulations, the method remained well-calibrated across simulated effect architectures. Application to Genebass summary statistics^19^ from 394,841 UK Biobank whole-exome sequences revealed widespread evidence of heterogeneous variant effects among genes exhibiting cross-phenotype burden signals. In *ALB*, for example, a subset of missense variants presented distinct effect patterns on albumin and calcium levels opposite to the direction of phenotype correlation. Protein structure-based analyses further showed enrichment of calcium-specific variants, as well as variants with opposing effects on albumin and calcium, near established calcium-binding regions of albumin, illustrating how allelic heterogeneity clarifies biological interpretation for gene-level cross-phenotype associations.

## Results

### Statistical test to model heterogeneous effects in genes with cross-phenotype associations

To characterize heterogeneity of variant effects within genes showing cross-phenotype associations, we developed ALLSPICE (ALLelic Spectrum of Pleiotropy Informed Correlated Effects), a likelihood-based statistical framework designed to distinguish differences in variant effect patterns within genes. ALLSPICE detects heterogeneous variant-level effects arising when distinct functional classes of variants generate different within-gene architectures of cross-phenotype burden associations. For example, predicted loss-of-function (pLoF) variants are expected to disrupt gene function through mechanisms such as nonsense-mediated decay, and therefore, often produce relatively concordant phenotypic effects across variants within a gene, whereas missense variants may alter protein function through diverse molecular mechanisms, potentially leading to greater heterogeneity in phenotypic effects across alleles.

We define an association as a significant burden signal between a gene-annotation pair and a phenotype (p_burden_ < 2.5 × 10^-6^; **Figure 1A**), and an association pair as a gene-annotation pair linked to two phenotypes through burden testing (**Figure 1B**). For each association pair, ALLSPICE evaluates whether rare protein-coding variants within the gene exhibit heterogeneous effects across the two phenotypes through a likelihood ratio test. In particular,

**Figure 1.**
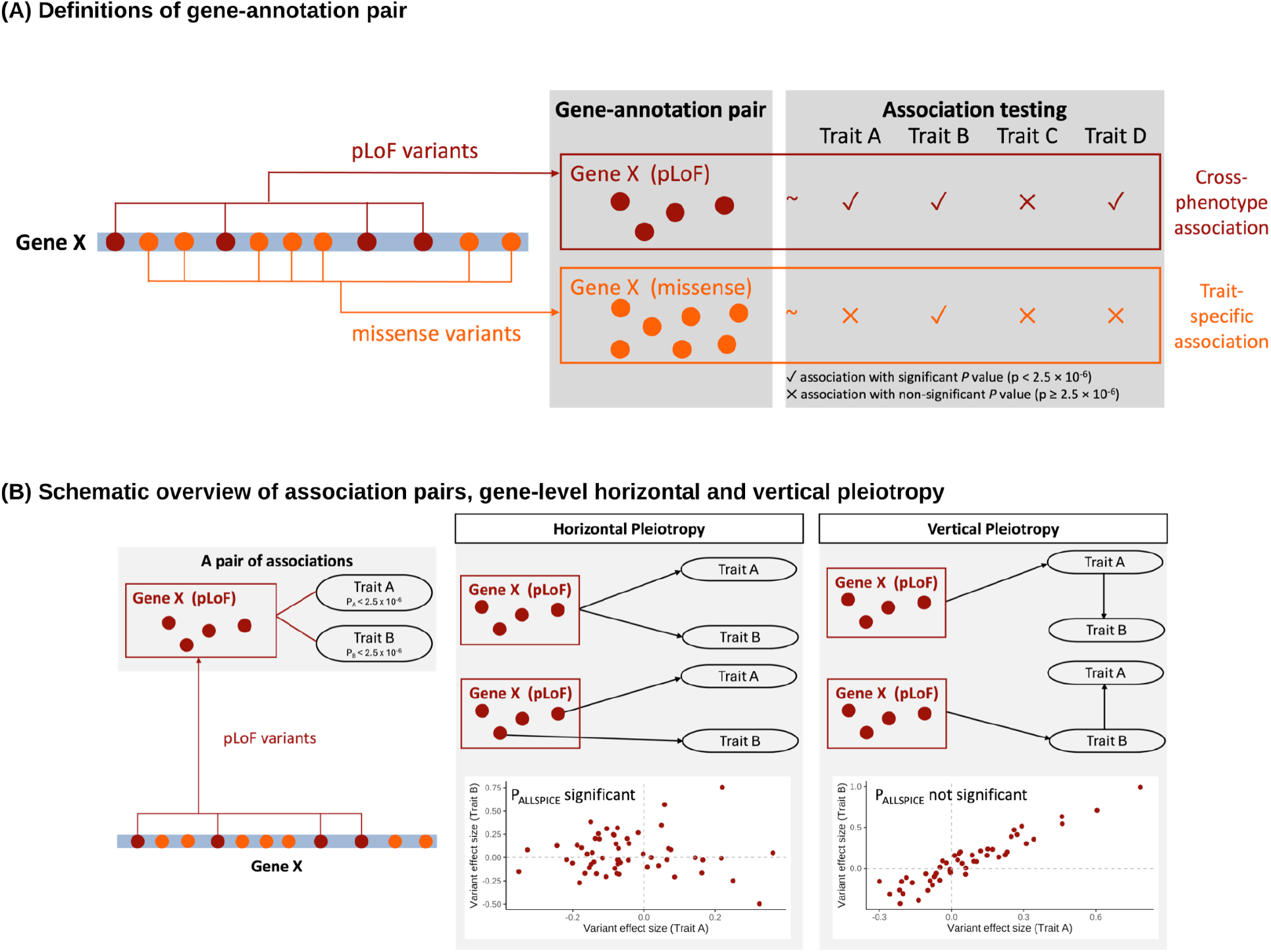
Definition of gene-annotation pairs and association pairs. **A**, Schematic representation of gene-annotation pairs, defined as sets of rare variants within a gene grouped according to functional annotations (pLoF, missense, synonymous, and pLoF+ missense) for burden testing, and of cross-phenotype associations, defined as a gene-annotation pair showing significant burden associations (p_burden_ < 2.5 × 10^-6^) with more than one phenotype. **B**, Definition of an association pair (left), schematic illustrations of gene-level horizontal pleiotropy (middle) and vertical (right) pleiotropy, and corresponding expected variant-level effect size patterns shown in the scatter plots.

ALLSPICE tests the null hypothesis *H*_0_: β_1_ = *c*β_2_, which assumes a perfectly proportional linear relationship between variant effect sizes for the two phenotypes, with the proportionality parameter *c* estimated by maximum likelihood under a closed-form linear regression framework that accounts for phenotypic correlation, and allele frequencies of rare variants within the same functional annotation class in a gene. The alternative hypothesis is *H*_1_: β_1_ ≠ *c*β_2_ for any value of *c*. ALLSPICE then constructs a likelihood ratio test statistic 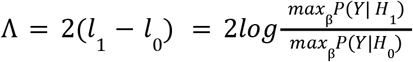 where *l*_1_ and *l*_0_ denote the maximized log-likelihoods under the alternative and null hypotheses, respectively. Because the likelihood is derived under a Gaussian linear model, the current implementation is restricted to continuous traits. Since ALLSPICE is designed for ultra-rare variants with allele frequency below 0.01%, linkage disequilibrium (LD) among analyzed alleles is assumed to be negligible. The significance of the statistical test is evaluated using the asymptotic chi-square distribution. A significant p-value indicates deviation from perfect proportionality, consistent with heterogeneous variant effects on the two phenotypes or, more generally, non-proportional allelic effects within the gene, whereas a non-significant p-value indicates that the observed variant effects are consistent with a shared linear relationship between the two phenotypes, conditional on their phenotypic correlation (**Methods**).

Although ALLSPICE does not explicitly classify pleiotropy subtypes, it is informative for distinguishing vertical pleiotropy from certain forms of horizontal pleiotropy: vertical pleiotropy is expected to yield proportional effects, whereas horizontal pleiotropy may produce heterogeneity when variants influence different molecular functions and downstream traits through distinct biological pathways^20^ (**Figure 1B**). In simulation studies, ALLSPICE presented uniformly distributed p-values under the null hypothesis (**Extended Data Figure 1** and **Figures S8-S11**), indicating appropriate type I error control and adequate statistical power under alternative models introducing allelic heterogeneity across phenotypes (**Extended Data Figure 2** and **Figures S12-S14**).

### Widespread cross-phenotype burden associations and allelic heterogeneity within genes

To assess the extent of cross-phenotype burden association in Genebass^19^, we curated 599 phenotypes with at least one significant gene-based rare variant burden signal (p_burden_ < 2.5 × 10^-6^; **Methods**), including 359 continuous traits and 240 binary traits. Further restricting to a maximal subset of 239 approximately independent phenotypes (r^2^ < 0.1; **Methods**), we identified 313 gene-annotation pairs associated with more than one phenotype, involving 185 unique genes (**SuppTable1**). Cross-phenotype burden signals were observed across all functional annotation classes, including 28% of pLoF, 23% of missense, 24 % of pLoF+missense, and 11% of synonymous gene-annotation pairs (**Figure 2A**). Across all annotation groups, the number of associated phenotypes per gene-annotation pair was overdispersed relative to both the Poisson expectation and the p-value-permuted null, indicating non-random structure beyond the conditional null expectations (**Table S1 and Figure S1**).

**Figure 2.**
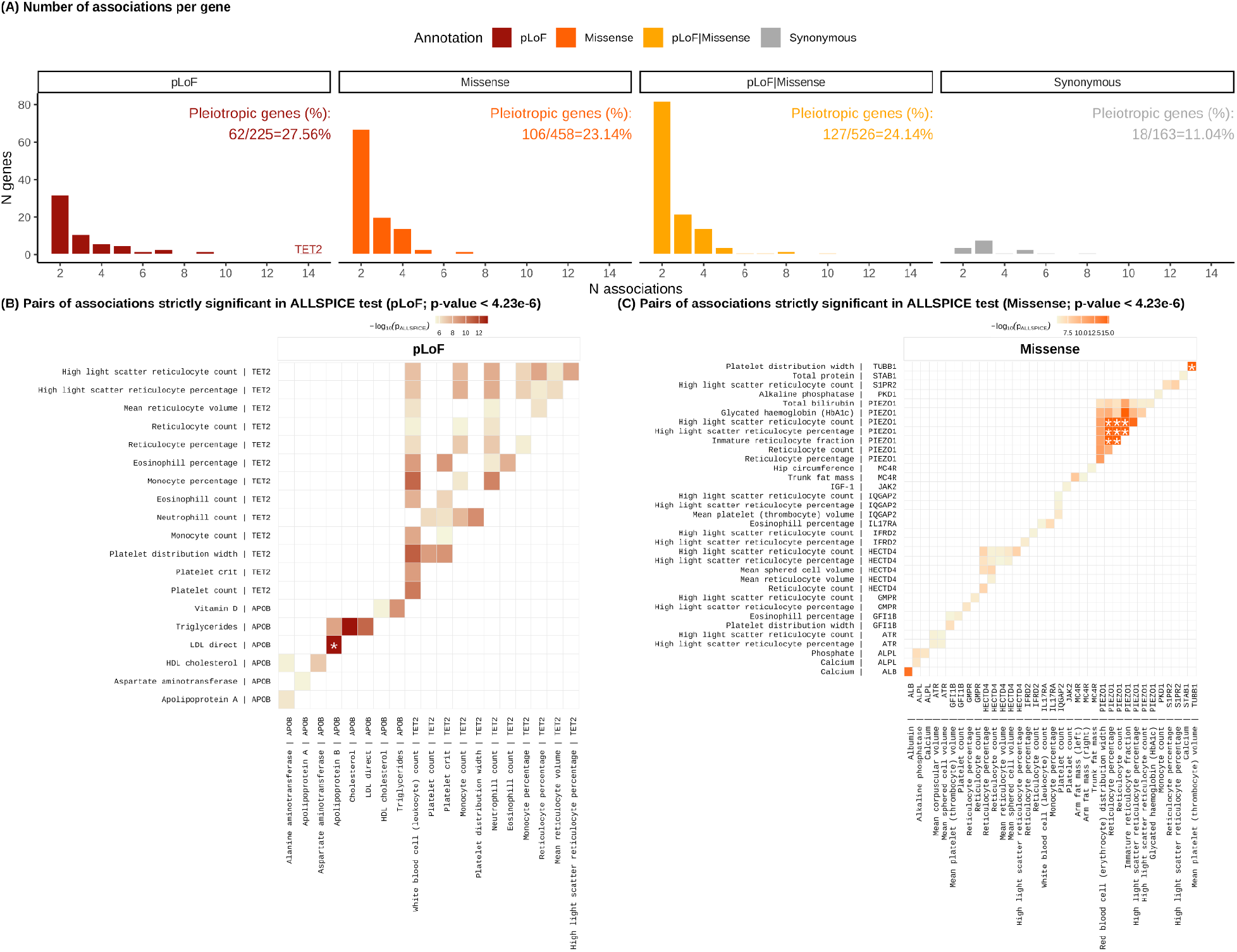
Summary of cross-phenotype associations and ALLSPICE results. **A**, Distributions of genes by the number of associated phenotypes (x-axis) across four functional annotation groups (panels and colors). Labels above the bars indicate the proportion of genes with more than one association among genes with at least one association in the corresponding annotation group. **B**, Heatmap of p_ALLSPICE_ for strictly significant pairs of pLoF associations: with phenotype pairs on the x- and y-axes, and genes labeled accordingly. Cell color indicates -log_10_(p_ALLSPICE_). Cells marked with asterisks denote association pairs with p_ALLSPICE_=0. **C**, Heatmap of p_ALLSPICE_ for strictly significant missense association pairs, using the same display format as in **B**.

To determine whether these cross-phenotype signals reflect homogeneous or heterogeneous allelic effects, we then assessed heterogeneity within a functional annotation class by applying ALLSPICE to 11,810 association pairs derived from the 359 continuous traits (4,708 pLoF, 5,911 missense, and 1,191 synonymous). Analyses were first restricted to gene-annotation pairs with at least two variants contributing to the corresponding burden tests (**SuppTable 3**). To further reduce potential confounding from linkage disequilibrium, we filtered rare protein-coding variants within each gene-annotation pair to those with MAF below 0.01%, and retained only association pairs with at least two variants after filtering. Among these 11,810 association pairs, 1,073 showed nominal evidence of heterogeneous variant effects (p_ALLSPICE_ < 0.05), including 292 pLoF, 681 missense, and 100 synonymous. After correction for multiple testing (p_ALLSPICE_ < 4.23 ×10^-6^; 0.05/11,810 tests), 124 association pairs remained significant (53 pLoF, 70 missense, and 1 synonymous; **Figure 2B-C**, and **Figure S15**). Consistent with power considerations, both the number of pLoF and the number of missense variants in a gene-annotation pair positively correlated with the likelihood for that association pair to be significant (correlation estimate = -0.09, p_pearson’s_=1.5 ×10^-21^ ; **Figure S15C**), indicating heterogeneous effects are more readily detected in genes with larger numbers of rare variants, whereas power is reduced for genes with sparse variant counts.

Notably, we identified 53 pairs of pLoF associations with significant ALLSPICE results (**Figure 2B**), despite pLoF variants often being assumed to have largely uniform effect directions. These significant association pairs were concentrated in two relatively long genes with broad biological relevance: *APOB* (CDS length 13,689; 10 significant association pairs) and *TET2* (CDS length 6006; 43 significant association pairs). *APOB* encodes a structural component of apoB-containing lipoproteins, and loss-of-function mutations in *APOB* cause familial hypobetalipoproteinemia, with reduced low-density lipoprotein cholesterol and apolipoprotein B^21^. *TET2* encodes an epigenetic regulator of hematopoietic stem and progenitor cells, and loss-of-function mutations in *TET2* affect myeloid differentiation and clonal expansion across blood cell lineages^22^. These well-characterized gene-specific effects provide context for the heterogeneous pLoF associations observed across phenotypes. We also applied ALLSPICE to the same set of association pairs restricted to ultra-rare protein-coding variants with Allele Counts (AC) < 5, which presented similar results consistent with those from rare protein-coding variants (AF < 0.0001) with a slightly refined set of significant signals (**Figure S18; SuppTable 6**).

### Examples of variant effect heterogeneity within genes

An illustrative example from ALLSPICE results is *ALB*, which encodes albumin and is associated with blood calcium and albumin levels through both pLoF (p_albumin_ = 2.40 × 10^-256^; p_calcium_ = 5.85 × 10^-40^) and missense (p_albumin_ = 1.73 × 10^-62^; p_calcium_ = 1.99 × 10^-16^) variants. Circulating calcium is partially albumin-bound; therefore, total serum calcium levels depend on albumin concentration^23,24^, as supported by the observed phenotypic correlation between albumin and calcium (r^2^ = 0.51). ALLSPICE result on this association pair, *ALB*-albumin and -calcium, reveals strong evidence of heterogeneous effects among rare missense variants (p_ALLSPICE_ = 5.5 × 10^-15^), whereas pLoF variants show no evidence of heterogeneity (p_ALLSPICE_ = 0.557), consistent with the uniform directional effects both observed and expected (**Figure 3A**). Notably, a missense variant at chr4:73412072:A:G (ENSP00000295897.4:p.Lys264Glu) is associated with increased calcium levels but decreased albumin levels, suggesting that the effect of some missense variants on calcium is not fully explained by their effect on albumin concentration.

**Figure 3.**
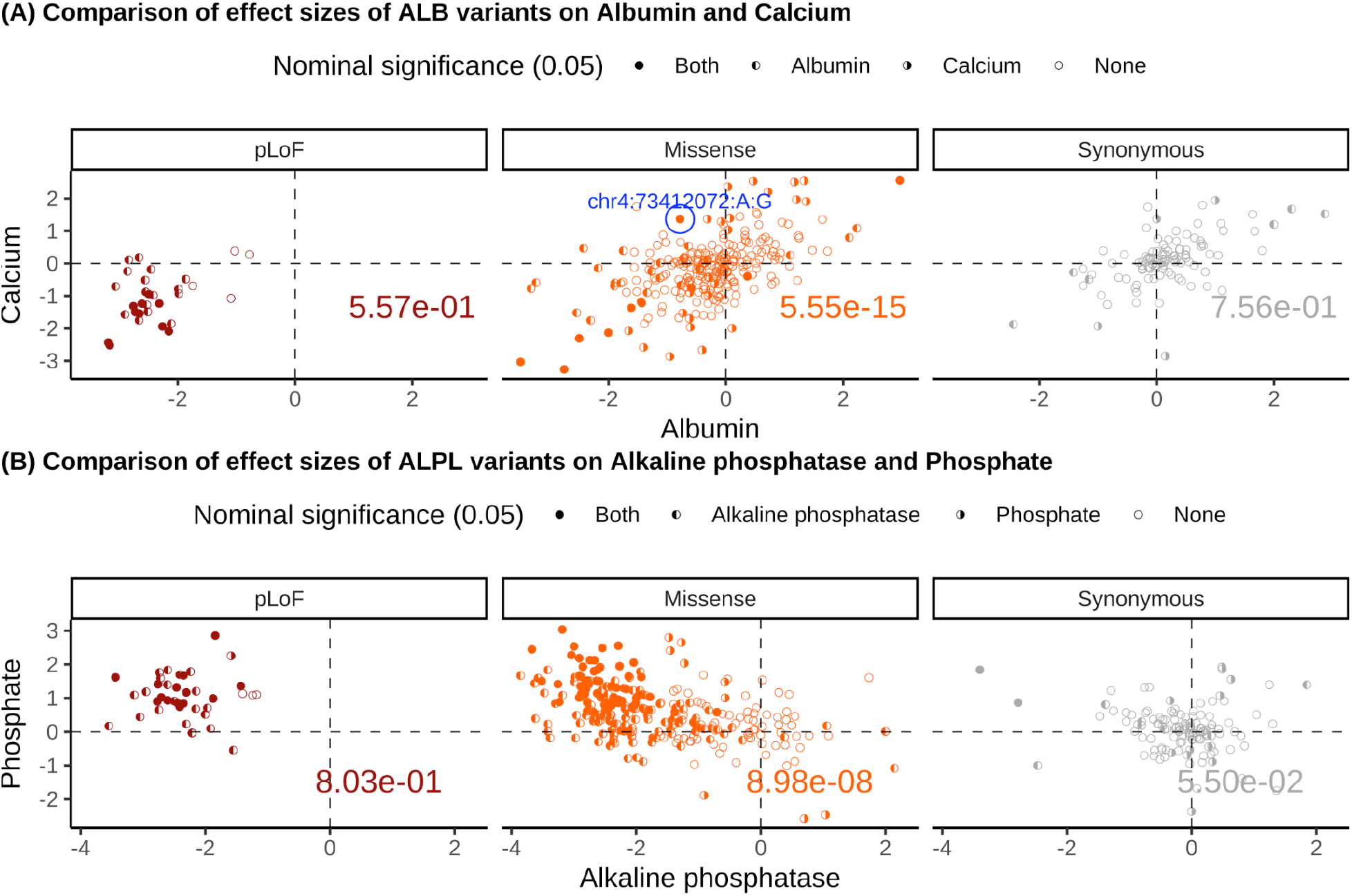
Examples of association pairs significant in ALLSPICE. **A**, Comparison of variant effect sizes on albumin (x-axis) and calcium (y-axis) among variants in *ALB*, stratified by functional annotation group (colors and columns). Point shapes indicate nominal significance (p_GWAS_ < 0.05) in single-variant association tests for the two phenotypes. The missense variant chr4:73412072:A:G, labeled and highlighted by a blue circle, is significantly associated with both traits but exhibits opposite directions of effect and represents one of the strongest contributors in the LOVO analysis. The value shown in the bottom right of each panel denotes p_ALLSPICE_ for the corresponding annotation group. **B**, Comparison of *ALPL* variant effect sizes on alkaline phosphatase (x-axis) to phosphate level (y-axis). Panels **A** and **B** share the same legend.

To assess which missense variants contribute most strongly to the ALLSPICE signal, we performed a leave-one-variant-out (LOVO) analysis, rerunning ALLSPICE repeatedly while removing one variant at a time and evaluating the changes in *c*_*MLE*_ and the p_ALLSPICE_. In the *ALB*-albumin and -calcium association pair, removal of 159 variants, primarily variants with near-zero effects on both phenotypes, increased ALLSPICE test significance (transparent points in **Figure S16A**), whereas removal of the remaining 235 variants attenuated the signal. Among these 253 variants, chr4:73412072:A:G produced the largest change in *c*_*MLE*_ (**Figure S16B; SuppTable 7**) and the 9th largest change in p_ALLSPICE_ (**Figure S16A; SuppTable 7**), indicating a disproportionate contribution of this variant to the observed ALLSPICE signal.

We further examined whether *ALB* missense variants associated with calcium versus albumin levels (p_burden_ < 0.05) show different spatial distributions on the *ALB* protein (**Methods**). Missense variants associated with calcium levels are located significantly closer to known calcium-binding sites in the 3D protein structure of *ALB*, compared with variants exclusively associated with albumin level (p_protein_ = 1.99 × 10^-10^; **Figure 4A, B**). Missense variants associated with albumin- and calcium-levels in an opposite direction also tend to be located closer to the calcium-binding sites at the 3D protein level (p_protein_ = 6.2 × 10^-4^; **Figure 4C, D**). These spatial patterns indicate that *ALB* missense variants near calcium-binding sites may affect calcium levels by directly disrupting ALB’s calcium-binding capacity, and thereby producing distinct functional effects on albumin-related downstream biological processes. The association pair *ALB*-albumin and -calcium also has a significant ALLSPICE result when restricted to ultra-rare missense variants with AC < 5 (p_ALLSPICE_ = 3.5 × 10^-7^; **Figure S18; SuppTable6**), for which the 3D protein distribution pattern persists (**Figure S19**).

**Figure 4.**
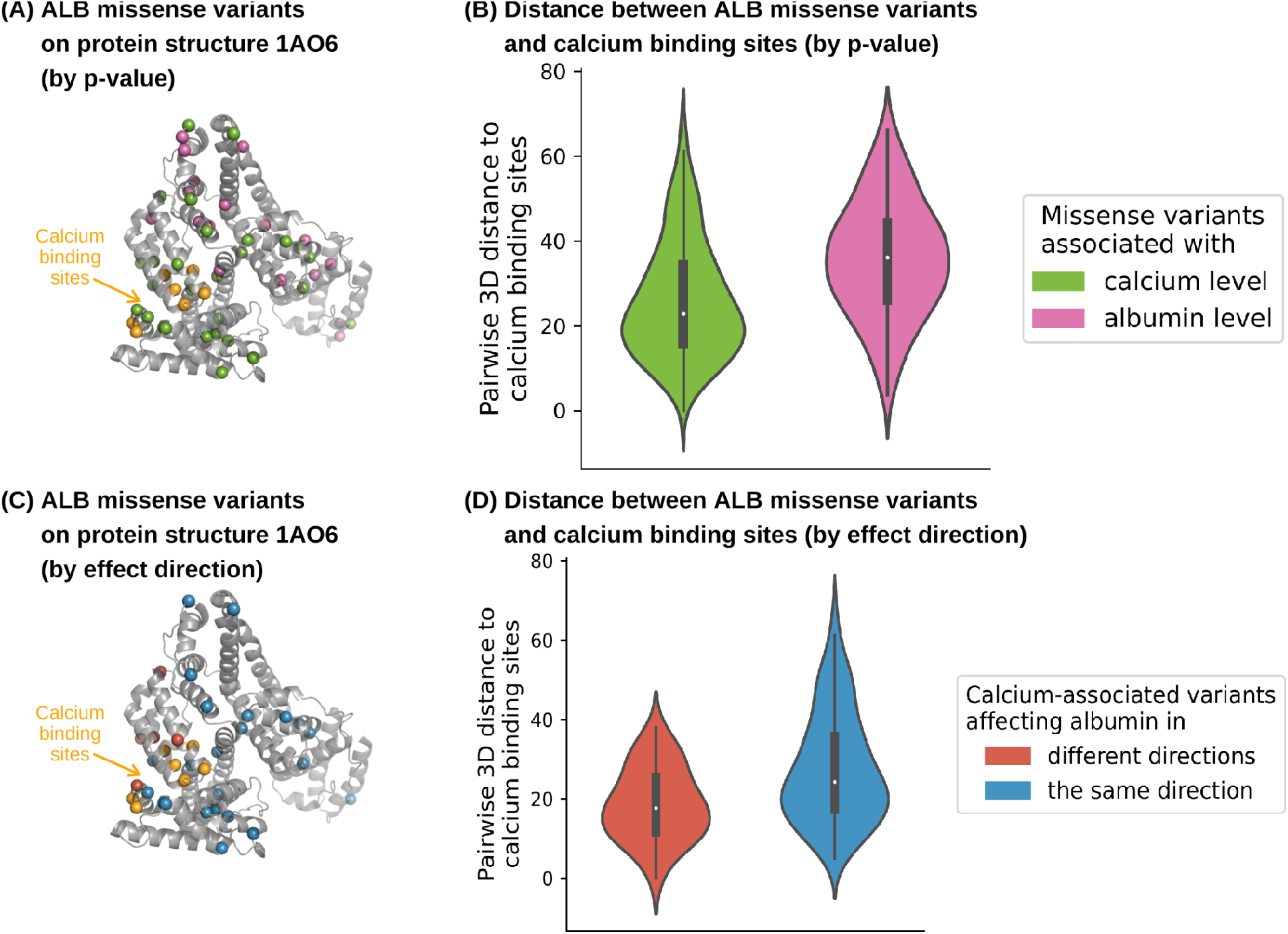
Rare protein-coding variants in *ALB* (AF < 0.0001) on ALB protein structure. **A**, Distribution of *ALB* missense variants associated with calcium level but not albumin level (pink), variants associated with albumin but not calcium level (green), and known calcium binding sites (orange) on the 3D structure of ALB protein (PDB ID: 1AO6). **B**, Comparison of pairwise distances from missense variants only associated with calcium (pink) or albumin-level (green) to calcium binding sites on ALB protein structure in the 3D space. Colors in panel **B** are defined the same as in panel **A. C**, Distribution of *ALB* missense variants associated with either albumin or calcium level that affect the two biomarker levels in the same (blue) or different (red) direction, and calcium binding sites (orange) on the 3D structure of ALB protein (PDB ID: 1AO6). **D**, Comparison of pairwise distances between missense variants to calcium binding sites on the ALB protein structure in the 3D space. Colors in panel **D** are defined the same as in panel **C**.

Another ALLSPICE-significant example is *ALPL*, which regulates alkaline phosphatase production and is associated with a wide range of biomarkers. Missense variants in *ALPL* are associated with calcium levels (p_burden_ = 4.15 × 10^-11^) and C-reactive protein (p_burden_ = 2.45 × 10^-6^; **Figure S17A**), while both pLoF and missense variants are associated with alkaline phosphatase and phosphate levels (pLoF: p_alkaline phospate_ = 1.99 × 10^-110^, p_phospate_ = 5.56 × 10^-8^; missense: p_alkaline phospate_ = 0, p_phospate_ = 2.17 × 10^-175^). Alkaline phosphatase catalyzes phosphate removal, leading to an inverse relationship between the two biomarkers. Consistent with this biological mechanism, most pLoF variants are associated with decreased alkaline phosphatase levels and increased phosphate levels, and show no evidence of heterogeneous variant effects across the two traits (p_ALLSPICE_ = 0.803), consistent with the role of pLoF variants in disrupting *ALPL* activity. In contrast, missense variants exhibit significant heterogeneity (p_ALLSPICE_ = 8.98 × 10^-8^; **Figure 3B**), displaying a mixture of opposing and concordant effects on alkaline phosphatase and phosphate, concordant with more heterogeneous functional consequences. Unlike *ALB*, missense variants in *ALPL* show no evidence of functional clustering in the 3D protein structure (**Figures S17B, C**).

## Discussion

In this study, we developed ALLSPICE, a statistical framework for detecting heterogeneous variant-level effects within genes exhibiting cross-phenotype associations using rare variant association summary statistics. By decomposing gene-level burden signals into variant-specific patterns across phenotypes, ALLSPICE identifies rare coding variants within the same gene that exert divergent effects while contributing to cross-phenotype gene-level associations. Application of ALLSPICE to rare variant association results for 359 continuous phenotypes in Genebass revealed substantial allelic heterogeneity underlying cross-phenotype burden signals, highlighting the value of ALLSPICE for dissecting rare variant pleiotropy beyond gene-level burden associations.

Complementary to Mendelian randomization, ALLSPICE characterizes heterogeneous variant effects within genes showing cross-phenotype associations rather than inferring causal relationships between traits under assumptions of instrument validity and predominantly vertical pleiotropy. Under a simple vertical pleiotropic model in which one trait mediates genetic effects on a second trait, variant effects are expected to remain approximately proportional across phenotypes. ALLSPICE therefore tests a null hypothesis of a linear relationship between variant effects across traits, where rejection indicates deviation from simple mediation, consistent with heterogeneous pleiotropic architecture, potentially reflecting horizontal pleiotropy or other sources of deviations from the model.

Several methodological limitations complicate the interpretation of rare variant pleiotropy. Because ALLPICE operates on rare variant association summary statistics, it inherits the same power limitations as rare variant association studies, including unstable effect estimates at low allele counts, unequal detectability across variant classes, and reduced sensitivity caused by annotation-dependent functional heterogeneity. These challenges are particularly relevant when comparing predicted loss-of-function and missense variants, which differ systematically in allele frequency, effect size, and annotation reliability. Interpretation is further complicated when apparent heterogeneity among predicted loss-of-function variants reflects misannotation, isoform-specific consequences, or assay-related artifacts rather than true biological divergence^4^. Similarly, nominal pleiotropic signals involving synonymous variants suggest that residual confounding, imperfect tagging of common variant effects, or technical artifacts in burden testing can contribute to shared associations, particularly at modest allele-frequency thresholds. Together, these limitations underscore the need for larger datasets, improved functional annotation, and more robust modeling of rare variant effects.

Beyond statistical power, limitations in medical ontology also affect the interpretation of cross-phenotype signals. Grouping phenotypes into broad domains and restricting to relatively independent phenotypes provides a practical framework for large-scale analysis, but remains an imperfect representation of biological and clinical relationships, often combining traits with distinct etiologies while separating phenotypes linked through shared pathophysiology. Consequently, correlated traits can generate apparent pleiotropy through overlapping clinical definitions, biomarker-disease relationships, or disease hierarchies. Although ALLSPICE accounts for phenotypic correlation within each tested association pair when evaluating variant-level heterogeneity, residual dependence arising from imperfect phenotype classification may still influence the interpretation of cross-phenotype associations.

From a methodological perspective, ALLSPICE is currently formulated for continuous traits only, and therefore, extending the framework to discrete outcomes and enabling joint comparison across continuous and discrete phenotypes remains an important direction for future development. ALLSPICE also relies on simplifying assumptions about rare variant association data, most notably negligible linkage disequilibrium(LD) among ultra-rare variants within genes. Although this assumption is often reasonable given the extremely low frequencies of such variants, within-gene LD can occur in practice and may inflate gene-based burden test statistics in proportion to the number of alleles per gene^25^. Individual-level genotype data would provide a more robust basis for directly modeling rare-variant correlation structure and improving calibration when LD is non-negligible. Relaxing these assumptions will require more flexible modeling approaches capable of accommodating more complex genetic architectures, particularly in genes with limited rare variants. Integrating complementary sources of genetic evidence, such as expression or protein quantitative trait loci, may further strengthen biological interpretation and help refine pleiotropic architectures identified by ALLSPICE.

## Methods

### Data description

We use gene-based burden test results from Genebass^19^, comprising 8,074,878 single-variant tests and 75,767 gene-based burden tests across 4,529 phenotypes (1,233 continuous and 3,296 binary), derived from whole exome sequencing data of 394,841 UK Biobank participants^26^. A total of 19,407 protein-coding genes were annotated with up to four functional groups: predicted LoF (pLoF), missense (including low-confidence pLoF variants and in-frame indels), synonymous, and non-synonymous (pLoF + missense). Variants included in burden tests were pre-filtered to those with minor allele frequency (MAF) < 0.01, and gene-based association testing was performed using SAIGE, as described in Genebass. Analyses were restricted to 58,559 high-quality gene-annotation pairs, following Genebass-recommended quality-control criteria, including a minimum of two variants per test, mean sequencing coverage ≥ 20, expected allele count (AC) ≥ 50, and genomic control inflation factor (λ_*GC*_) > 0.75 for synonymous tests. Expected AC was defined as the combined allele frequency (CAF) of the gene-annotation group multiplied by the number of cases for the corresponding phenotype.

### Phenotype curation

Starting from the 4,529 phenotypes available in Genebass, we focus on biological measurements and health-related outcomes by excluding phenotypes related to hospital operations (1,704), treatment/medications (435), customized phenotypes (177), and non-endpoint phenotypes (64), including injuries and age-related measures. Of the remaining 2,149 phenotypes after the initial curation, we further excluded 1,550 phenotypes with no gene associations reaching exome-wide significance (p_burden_ < 2.5×10^-6^). This filtering results in a final set of 599 phenotypes with at least one gene-based association. To facilitate analyses of cross-phenotype associations across largely independent phenotypes, we additionally derived a subset of 239 non-redundant phenotypes with pairwise phenotypic correlation below 0.1.

### Statistical test for variant effect heterogeneity between continuous traits

When a gene is associated with multiple phenotypes, an important question is whether the same causal variants underlie these associations. We introduce ALLSPICE, a likelihood ratio test to identify variant effect heterogeneity within genes showing cross-phenotype associations, defined as non-proportional variant-level effects across two traits. The likelihood ratio test explicitly accounts for uncertainty in effect size estimates arising from sampling variation, which influences the shape of the likelihood function through its dependence on both sample size and allele frequency. Under the null hypothesis of perfectly proportional effects, the likelihood-ratio statistic follows an asymptotic chi-square distribution, with the corresponding critical values reflecting the variation expected from finite-sample noise. ALLSPICE operates efficiently on summary statistics from single-variant association tests, accounts for phenotypic correlation between traits, and does not require individual-level genotype or phenotype data.

We assume that *n* samples are sequenced for a gene with *m* variants. Given a pair of phenotypes, let *y*_*i*_ = (*y*_*i*1_,⋯, *y*_*in*_)^*T*^ be the *i*_*th*_ (*i* = 1, 2) phenotype for the *n* samples and β_*i*_ = (β_*i*1_,⋯, β_*im*_)^*T*^ be the effect sizes of *m* variants on the *i*_*th*_ phenotype. For single-variant association summary statistics on continuous phenotypes, we consider the linear regression model *Y* = β*X* + ϵ, where *Y* = (*y*_1_, *y*_2_)^*T*^, β = (β_1_, β_2_)^*T*^, where *X* is an *m* × *n* matrix denoting the genotypes of *n* samples at *m* variation sites, and ϵ_2×*n*_ ∼ *N*(0, Δ^2^) is the environmental noise term. We then define *Y* ∼ *N*(β*X, R*), where 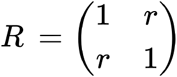 with *r* representing the phenotypic correlation between the pair of phenotypes, which is empirically computed as the correlation between effect sizes of all synonymous variants across the exome on the two phenotypes. We restrict to only the rare protein-coding variants with minor allele frequency < 0.01%, limiting the effect of linkage disequilibrium (LD), and assume that the variants are independent, as no individuals have two or more variants among the *m* variants:

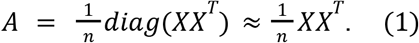

We test the null hypothesis:

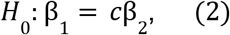

for some constant *c*, which describes the case when the effect sizes of individual variants within a gene for the pair of phenotypes are perfectly proportional to each other. The test statistic is defined as

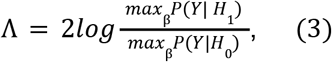

following a chi-square distribution with *m* − 1 degrees of freedom.

First, we consider the simplest version, where *m* = 1 and *n* = 1. For the *k*_*th*_ individual (*k* = 1) and the *j*_*th*_ variant (*j* = 1), we have

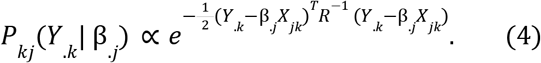

Denoting 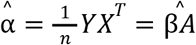, then we have

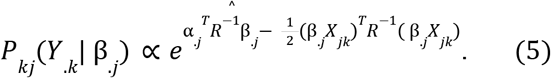

Therefore, for *n* unrelated samples and one variant, we have

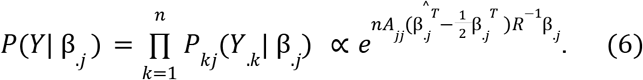

We then adapt the likelihood to the null and alternative hypotheses and compute the maximum likelihood estimator (MLE) of true effect sizes β_1_ and β_2_ under both conditions. The MLEs of effect size under the null are

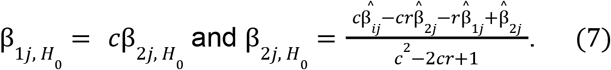

And those for the alternative hypothesis are simply the summary statistics

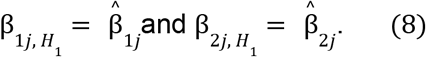

Therefore, by plugging the MLEs back into the likelihood ratio, we have

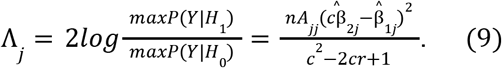

The final test statistic is computed as

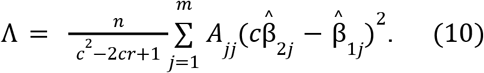

To formulate the unknown parameter *c*, we take the derivative of Λ(*c*)and compute the two extrema of Λ, which are

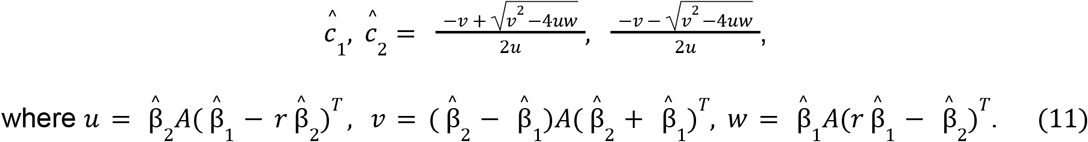

To obtain the maximum likelihood of the null hypothesis, we select the extremum that minimizes the value of Λ, which is ĉ = *max* (ĉ_1_, ĉ_2_) when *u* > 0 and ĉ = *min*(ĉ_1_,ĉ_2_) when *u* < 0. Finally, we have the equation for the likelihood ratio test statistic

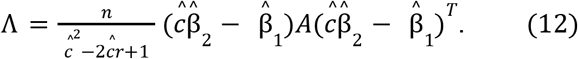

When phenotypic correlation *r* between two traits approaches one, variant effect sizes on the two phenotypes become nearly identical, leading the estimated slope ĉ to be close to one and thus the denominator of the test statistic to be close to zero (Equation 10). We therefore recommend avoiding the application of ALLSPICE to pairs of associations involving highly positively correlated phenotypes. This is also consistent with the intended use of ALLSPICE, which is designed to highlight variants with heterogeneous effects across traits.

### Simulation design

To evaluate the performance of ALLSPICE, we conducted extensive simulations across a wide range of parameter combinations. We simulated data under 1,584 parameter configurations under the null hypothesis and 792 scenarios under the alternative hypothesis to assess statistical power.

Given the definitions of null and alternative hypotheses, the simulation differs in how effect size vectors are generated. Under the null hypothesis, the true effect sizes for the second phenotype (β_2_) were sampled from a mixture distribution β_2_ = *g*_2_ *z*_2_, *z*_2_ ∼ *N*(0, *σ*^2^) where *g*_2_ ∼*Bernoulli*(π) indicates whether an individual variant has a nonzero effect and π denotes the probability of those variants with nonzero effects. Effect sizes for the first phenotype (β_1_) were then defined as β_1_ = *c*β_2_, enforcing perfect proportionality between effects across the two phenotypes.

Under the alternative hypothesis, we consider three distinct scenarios in which the perfect proportional linearity is violated.

1. Independent effects: effect sizes for the first phenotypes β_1_ were generated independently using the same mixture model as for the second phenotype β_2_, representing completely uncorrelated effects across phenotypes.
  a. β_1_=*g*_1_*z*_1_ with *g*_1_∼*Bernoulli*(π) and *z*_1_∼*N*(0, σ^2^)
  b. β_2_=*g*_2_*z*_2_ with *g*_2_∼*Bernoulli*(π) and *z*_2_∼*N*(0, σ^2^)
2. Correlated but non-proportional effects: β_1_ was generated to have a moderate Pearson correlation (ρ = 0. 5) with β_2_, to form a scenario different from the null hypothesis. Specifically, we first sampled two latent variables *z*_1_, *z*_2_ ∼ *N*(0, *σ*^2^), and defined 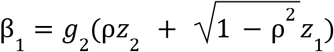, while keeping β_2_ = *g*_2_ *z*_2_. This construction preserves equal marginal variances while inducing correlated but not proportional effects across the two phenotypes.
3. Non-linear effects: to model deviations from linearity, effect sizes for the first phenotype β_2_ were generated as a nonlinear function of β_2_, β_1_ = *g*_1_ (β_2_ ^2^+ β_2_ + ϵ), ϵ ∼ *N*(0, *σ*^2^), where *g*_1_ ∼*Bernoulli*(π) and β_2_ is defined the same as previous scenarios. This scenario introduces nonlinear dependence and additional noise, representing complex genetic architectures in which variant effects on one phenotype depend nonlinearly on their effects on another phenotype.

We fixed the number of samples *n* at 1,000, and then varied parameters over the ranges as specified below, performing 100 simulations for each parameter combination:

- Number of variants within a gene *m* ∈ {5, 10, 20, 100},
- Slope between variant-level effect sizes on two phenotypes

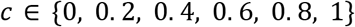
- Phenotypic correlation between the two phenotypes

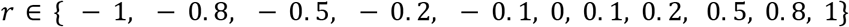
- Variance of the component normal distribution of effect sizes *σ*^2^ ∈ {0. 01, 0. 1, 1}
- Weight for the mixture distribution of variant effects π ∈ {0. 5, 0. 8}

We simulated minor allele counts for *m* rare protein-coding variants from a *Uniform*(1, 100) distribution, and the genotype matrix *X*_*m*×*n*_ as a binary (0, 1) matrix, where each variant *j* contains *nA*_*jj*_ minor alleles randomly distributed across samples. Phenotype values for the two traits were generated from a multivariate normal distribution *N*(β*X, R*), which encodes the phenotypic correlation between the traits. For each simulation, allele counts *A*, effect sizes β, and genotype matrix *X* were independently randomized, since fixing for any of these components did not substantially affect the test statistics. Summary statistics were estimated as 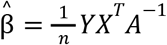, which approximates the effect sizes under a linear regression model. Finally, for the null hypothesis, we computed the maximum likelihood estimate of *c* by maximizing the likelihood under the null, thereby minimizing the test statistic.

### Simulation results

We applied ALLSPICE to simulated data across all combinations of parameters. Under the null hypothesis with *σ*^2^ = 1 and π = 0. 5, the distribution of p_ALLSPICE_ was calibrated across all scenarios (**Extended Data Figure 1A; SuppTable 9**). We computed the empirical type I error rate as the proportion of tests rejecting the null hypothesis at the nominal significance level (p_ALLSPICE_ < 0.05). Across most scenarios, type I error rates were centered around 0.05. Although error rates increased slightly with the number of variants, they remained well controlled below 0.1 (**Extended Data Figure 1B**). When *σ*^2^ was reduced to 0.01 or 0.1, the test exhibited mildly conservative patterns compared to *σ*^2^ = 1, reflecting the smaller effect sizes simulated as *σ*^2^ decreases (**Figures S8, S9; SuppTable 10, 11**). Results for higher mixture proportions (π = 0. 8) were similarly well calibrated compared to π = 0. 5 (**Figures S10-S12; SuppTable 9-11**).

Empirical statistical power was computed as the proportion of the tests rejecting the null hypothesis at a nominal significance level of 0.05 under the alternative. Parameter configuration is the same as previously described for simulations under the null scenario. We fixed the sample size *n* at 1,000 and *σ*^2^ at 1, and performed 1,000 simulations for each parameter configuration. Statistical power increased with the number of variants included in the gene, exceeding 0.9 when at least 10 variants were present, which is consistent across three alternative scenarios (**Extended Data Figure 2; SuppTable 12, 15**, and **18**). However, when *σ*^2^ was reduced to 0.01 or 0.1, the overall scale of effect sizes for both β_1_ and β_2_ decreased, thus leading to a reduction in power to detect significant association pairs, particularly for the scenarios with independent effect sizes (**Figure S13 and Figure S14; SuppTable 13, 14, 16, 17, 19**, and **20**).

### Protein structure-based analysis

Pairwise distances between residuals on the 3D protein structures were calculated as the Euclidean distances between the alpha carbons of the corresponding residues, using their atomic coordinates from the Protein Data Bank (PDB). p-values (p_protein_) indicating the differences in distance distributions between groups were computed through a two-tailed Mann-Whitney U test (**Figure 4, Figure S17**, and **Figure S19**).

## Extended Data Figures

**Extended Data Figure 1.**
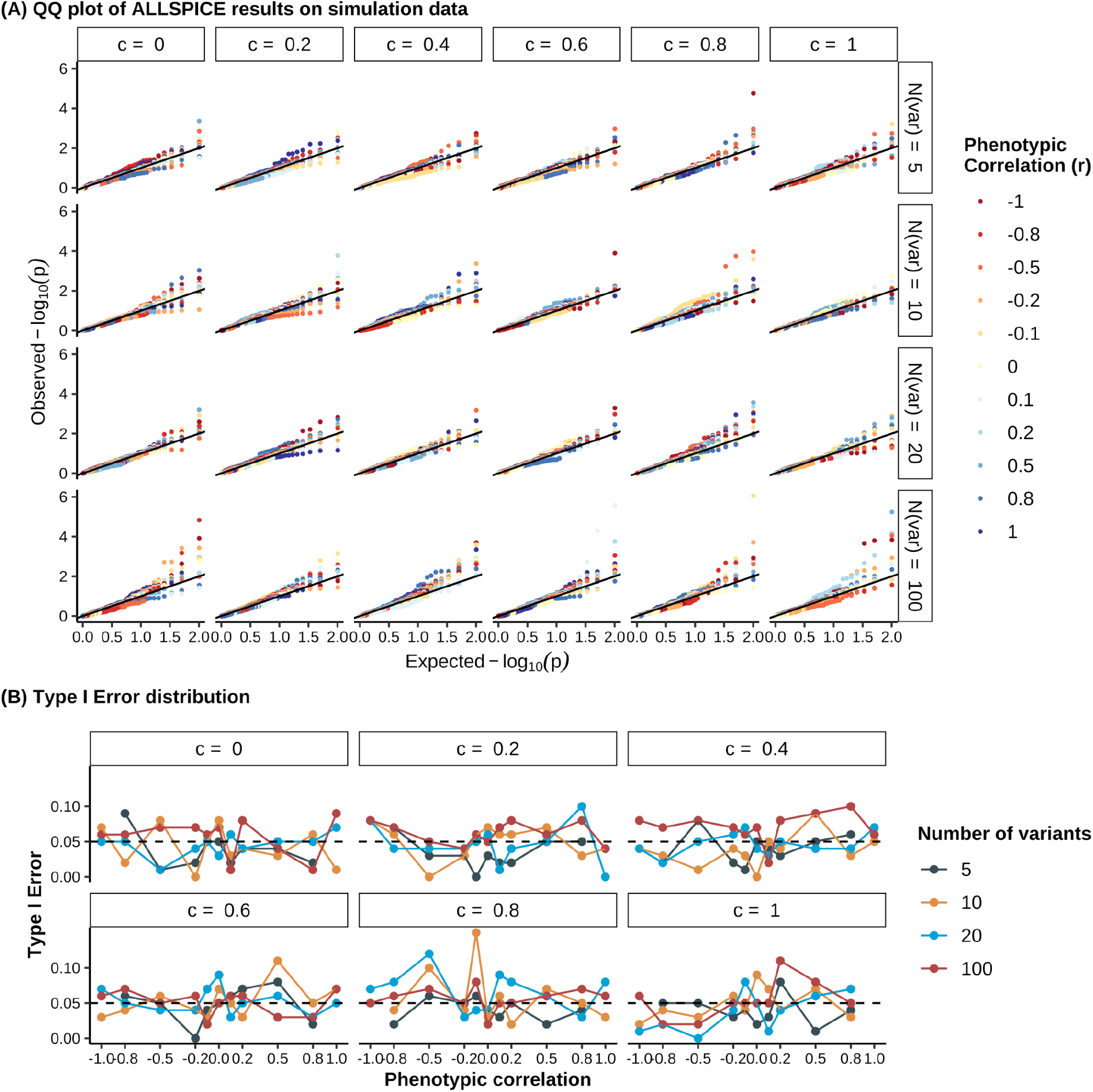
ALLSPICE test results on data simulated under the null hypothesis *H*_0_: β_1_ = *c*β_2_ across 264 scenarios when the variance *σ*^2^= 1 and the mixture component weight π = 0. 5. **A**, QQ plot of test results from the simulated null data described above. The solid black line denotes the line y=x. Rows correspond to the number of variants included, and the columns indicate the true value of the slope parameter *c* relating the two effect size vectors β_1_, β_2_ used in the simulation. **B**, Distribution of type I error rates (y-axis) under the null hypothesis *H*_0_: β_1_ = *c*β_2_ across combinations of phenotypic correlation (x-axis), number of variants (colors), and true values of slope *c* (panels). The dashed horizontal line represents the nominal significance level y = 0.05.

**Extended Data Figure 2.**
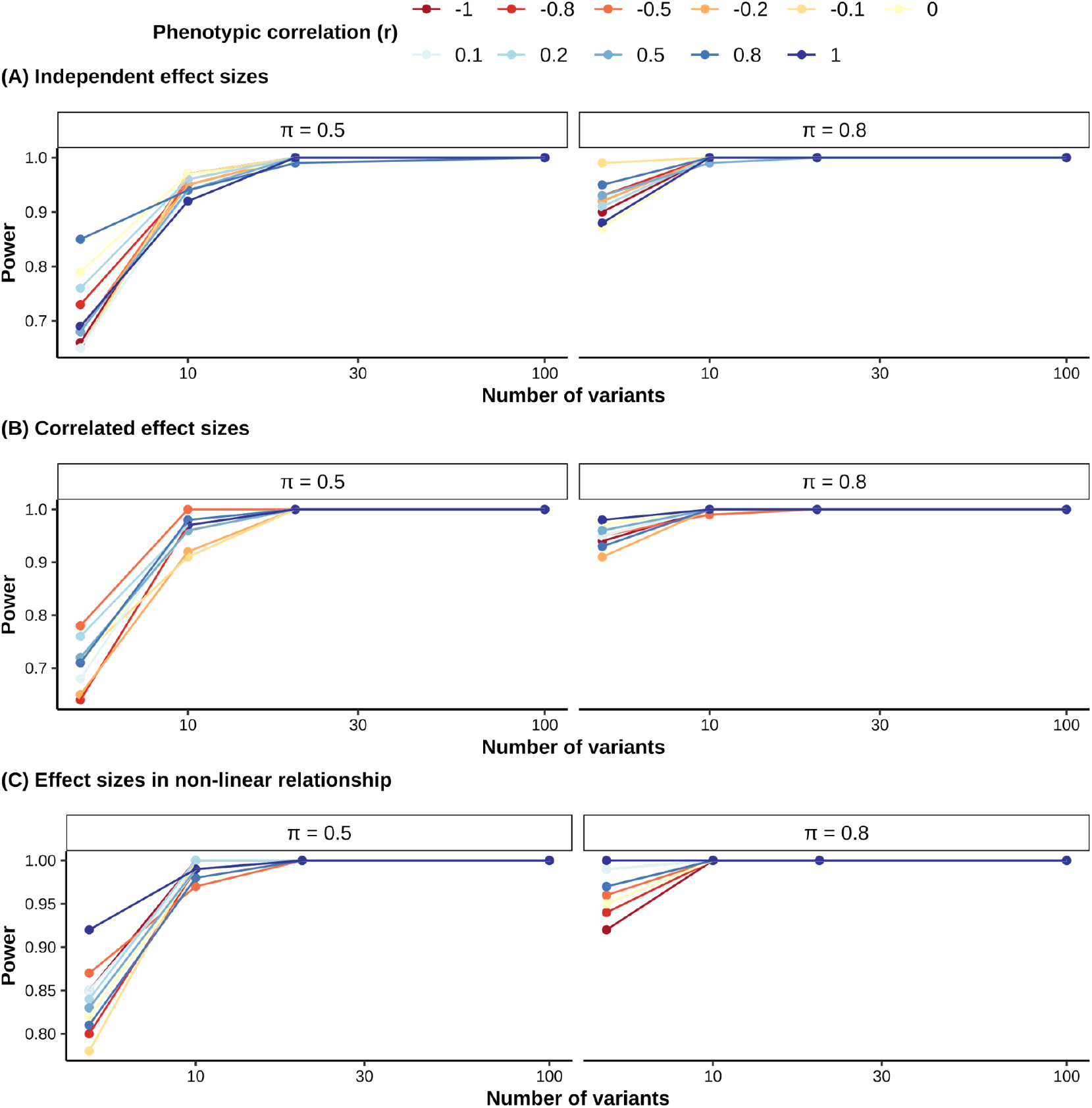
Distribution of empirical statistical power of ALLSPICE test under the alternative hypotheses (*σ*^2^ = 1). Empirical power (y-axis) was evaluated in simulated data across phenotypic correlation (color), number of variants (x-axis), and mixture component weights π (columns), indicating the probability that a variant effect is drawn from the non-zero component, modeled as a standard normal distribution *N*(0, *σ*^2^), where *σ*^2^ = 1.Results are shown for three alternative scenarios: **A**, independent effect sizes across phenotypes; **B**, correlated but non-proportional effect sizes; **C**, non-linear relationship between effect sizes.

## Resources

GitHub repository: https://github.com/atgu/ALLSPICER

## Acknowledgements

Grant from the Swiss National Science Foundation to C.A. (P500-3_235131) D.P. is supported by the Netherlands Organization for Scientific Research - Gravitation project ‘BRAINSCAPES: A Roadmap from Neurogenetics to Neurobiology’ (024.004.012)

## References

1. Watanabe, K. et al. A global overview of pleiotropy and genetic architecture in complex traits. Nat. Genet. 51, 1339–1348 (2019).

2. Sivakumaran, S. et al. Abundant pleiotropy in human complex diseases and traits. Am. J. Hum. Genet. 89, 607–618 (2011).

3. Levin, M. G. et al. Genome-Wide Assessment of Pleiotropy Across >1000 Traits from Global Biobanks. medRxiv (2025) doi:10.1101/2025.04.18.25326074.

4. Solovieff, N., Cotsapas, C., Lee, P. H., Purcell, S. M. & Smoller, J. W. Pleiotropy in complex traits: challenges and strategies. Nat. Rev. Genet. 14, 483–495 (2013).

5. Davey Smith, G. & Hemani, G. Mendelian randomization: genetic anchors for causal inference in epidemiological studies. Hum Mol Genet 23, R89–98 (2014).

6. O’Connor, L. J. & Price, A. L. Distinguishing genetic correlation from causation across 52 diseases and complex traits. Nat. Genet. 50, 1728–1734 (2018).

7. Valverde, P., Healy, E., Jackson, I., Rees, J. L. & Thody, A. J. Variants of the melanocyte–stimulating hormone receptor gene are associated with red hair and fair skin in humans. Nature Genetics 11, 328–330 (1995).

8. Lazarev, D., Chau, G., Bloemendal, A., Churchhouse, C. & Neale, B. M. GUIDE deconstructs genetic architectures using association studies. bioRxiv (2025) doi:10.1101/2024.05.03.592285.

9. Liang, X., Sha, Q., Rho, Y. & Zhang, S. A hierarchical clustering method for dimension reduction in joint analysis of multiple phenotypes. Genet Epidemiol 42, 344–353 (2018).

10. Bulik-Sullivan, B. et al. An atlas of genetic correlations across human diseases and traits. Nat. Genet. 47, 1236–1241 (2015).

11. Jee, Y. H. et al. Dissecting pleiotropy to gain mechanistic insights into human disease. Nature Reviews Genetics 27, 292–305 (2025).

12. Shi, H., Mancuso, N., Spendlove, S. & Pasaniuc, B. Local Genetic Correlation Gives Insights into the Shared Genetic Architecture of Complex Traits. Am. J. Hum. Genet. 101, 737–751 (2017).

13. Ballard, J. L. & O’Connor, L. J. Shared components of heritability across genetically correlated traits. Am. J. Hum. Genet. 109, 989–1006 (2022).

14. Wu, M. C. et al. Rare-variant association testing for sequencing data with the sequence kernel association test. Am. J. Hum. Genet. 89, 82–93 (2011).

15. Sinnott-Armstrong, N. et al. Genetics of 35 blood and urine biomarkers in the UK Biobank. Nat. Genet. 53, 185–194 (2021).

16. Chami, N. et al. Exome Genotyping Identifies Pleiotropic Variants Associated with Red Blood Cell Traits. Am. J. Hum. Genet. 99, 8–21 (2016).

17. Satterstrom, F. K. et al. Autism spectrum disorder and attention deficit hyperactivity disorder have a similar burden of rare protein-truncating variants. Nat. Neurosci. 22, 1961–1965 (2019).

18. Collaborative, E. Exome sequencing of 20,979 individuals with epilepsy reveals shared and distinct ultra-rare genetic risk across disorder subtypes. Nature Neuroscience 27, 1864–1879 (2024).

19. Karczewski, K. J. et al. Systematic single-variant and gene-based association testing of thousands of phenotypes in 394,841 UK Biobank exomes. Cell Genom 2, 100168 (2022).

20. Laddach, A., Ng, J. C. F. & Fraternali, F. Pathogenic missense protein variants affect different functional pathways and proteomic features than healthy population variants. PLoS Biol. 19, e3001207 (2021).

21. Peloso, G. M. et al. Rare Protein-Truncating Variants in APOB, Lower Low-Density Lipoprotein Cholesterol, and Protection Against Coronary Heart Disease. Circ Genom Precis Med 12, e002376 (2019).

22. Guarnera, L. & Jha, B. K. TET2 mutation as prototypic clonal hematopoiesis lesion. Semin Hematol 61, 51–60 (2024).

23. Desgagnés, N., King, J. A., Kline, G. A., Seiden-Long, I. & Leung, A. A. Use of Albumin-Adjusted Calcium Measurements in Clinical Practice. JAMA Netw Open 8, e2455251–e2455251 (2025).

24. Besarab, A., DeGuzman, A. & Swanson, J. W. Effect of albumin and free calcium concentrations on calcium binding in vitro. J Clin Pathol 34, 1361–1367 (1981).

25. Weiner, D. J. et al. Polygenic architecture of rare coding variation across 394,783 exomes. Nature 614, 492–499 (2023).

26. Bycroft, C. et al. The UK Biobank resource with deep phenotyping and genomic data. Nature 562, 203–209 (2018).

