## supplementary_information for "Effect heterogeneity reveals complex pleiotropic effects of rare coding variants"


**Permutation and Poisson dispersion test** **6**

Figure S1 | Permutation results. A, Distribution of the proportion of genes with cross-phenotype associations (x-axis) across permutations for each functional annotation group (columns and colors). B, Distribution of the proportion of genes with cross-phenotype associations across permutations, stratified by functional annotation group (columns and colors) and cumulative allele frequency interval (rows). The vertical dashed line in both A and B denotes the observed proportion in real gene-level association data. 7

Table S1 | Poisson dispersion test for the number of associations per gene-annotation pair. The test evaluates whether the observed distribution of association counts per gene is consistent with a Poisson model under the assumption of independent associations. A significant p-value here indicates overdispersion relative to the Poisson expectation, reflecting an excess of cross-phenotype relative to what would be expected if trait associations were independent. The $\chi$2 statistic is computed as i=1Ngene(Xi-X)2 X within each functional annotation group, where X denotes the mean association count across genes, with the number of associations being Xi. 8

**Pervasive extent of gene-level cross-phenotype associations** **9**

Figure S2 | Proportion (dot and lines; left y-axis) and count (bars; right y-axis) of genes with single-trait and cross-trait associations (rows) among 239 approximately independent traits (r2 < 0.1), across four CAF intervals (x-axis), stratified by functional annotation group (columns and colors). Proportions are calculated relative to the total number of genes is the total number of genes within each category. 10

Table S2 | Logistic regression results assessing the effects of CAF and CDS length on association status. Analyses were performed among genes with at least one association, and among all genes, including those with no associations, stratified by functional annotation group. 11

Figure S3 | Gene-level cross-phenotype associations across all 37 functional gene categories. For each category and functional annotation group, the proportion of genes with cross-phenotype associations is computed as the number of genes with two or more associations divided by the number of genes with at least one association (SuppTable3). Transparent points indicate categories with fewer than 5 genes tested. The dashed line denotes the corresponding proportion computed across all genes. Error bars represent standard errors. A, Associations across the 599 curated phenotypes; B, associations across the 239 approximately independent phenotypes (r2 < 0.1). 13

Figure S4 | Number of protein-protein interactions across association categories. The mean number of protein-protein interactions (y-axis) is shown for genes associated with no association, a single association, or multiple associations (x-axis), stratified by functional annotation group (colors and columns). Numbers above points indicate the total number of genes in each group, and error bars denote 95% confidence intervals. A, Associations across the 599 curated phenotypes; B, associations across the 239 approximately independent phenotypes (r2 < 0.1). 15

Pleiotropy manifests in diverse forms: allelic series and domain-level associations 15

Table S3 | Genes for which pLoF variants are associated with disease phenotypes and missense variants with biomarkers and no diseases, or vice versa. Disease phenotypes are shown in bold. 17

Table S4 | Gene burden associations (pburden < 2.5 × 10-6) of pLoF variants in GIGYF1 and KDM5B across phenotypic domains. 19

Figure S5 | Phenotypic correlation matrices for phenotypes listed in Table 1. Matrices are shown for phenotypes associated with A, ATM; B, PKD1; C, TMPRSS6; D, JAK2; E, SLC34A3; F, FAM234A; and G, MICA. Variant annotation groups are indicated alongside the phenotype labels on both the x- and y-axes. 19

**Genes with cross-domain associations across phenotypic and disease domains 20**

Figure S6 | Pervasive gene-level cross-domain associations across six broader phenotypic domains: A, Number of phenotypes in each of the six phenotypic domains (color-coded); B, Proportion of genes with trait-specific, domain-specific, and multi-domain associations defined at the phenotypic domain level, among gene-annotation pairs with at least one association across functional annotation groups (color-coded). The blue bar shows the common variant results from Watanabe et al. 20191; C, Number of genes associated with each combination of phenotypic domains. The bars indicate the number of genes with cross-domain associations, as shown in the bottom matrix, where each column represents a domain combination that shares at least one gene-level association. Colors are consistent between panel A and the bottom matrix in panel C. D, Proportion of constrained genes, defined as those in the lowest LOEUF decile (y-axis), with no association, trait-specific, domain-specific, or multi-domain associations (x-axis) across functional annotation groups (color and column). Error bars represent 95% confidence intervals. 21

Figure S7 | Gene-level cross-domain associations across 17 disease sub-domains. A, Number of diseases in each of the 17 disease sub-domains; B, Proportion of genes with trait-specific, domain-specific, and multi-domain associations at the disease sub-domain level, among gene-annotation pairs with at least one association across, stratified by functional annotation group. C, Number of genes associated with each combination of disease sub-domains. Each column of the bottom matrix represents a distinct sub-domain combination sharing gene-level associations. Bars indicate the number of genes with cross-disease-domain associations defined by the corresponding dot combinations in the matrix below. Colors are consistent between panel A and the matrix in panel C. 22

Table S5 | Comparison with common variant results from Watanabe et al. (2019). Shown are the proportion and number of genes with cross-phenotype, multi-domain, domain-specific, and trait-specific associations across six broad phenotypic domains (599 phenotypes), stratified by functional annotation group. 24

Table S6 | Proportion of genes with cross-phenotype, multi-domain, domain-specific, and trait-specific associations across 17 disease sub-domains, stratified by functional annotation group. 25

Table S7 | Correlations of the number of associations per gene across phenotypic domains, the values in the upper triangle are correlations between the number of associations per gene within each pair of domains, the values in the lower triangle are correlations between the binary vector, whether there is any association per gene within each pair of domains. 25

Table S8 | Logistic regression results for the impact of constraint level and gene CDS length on the pLoF group of a gene having multi-domain, domain-specific, and trait-specific associations compared to having no association. 26

**ALLSPICE testing on simulated data 27**

Figure S8 | ALLSPICE test results on data simulated under the null hypothesis H0:1=c2 across 264 scenarios when 2=0.1 and =0.5. A, QQ-plot of test results on the simulated data as described above. The solid black line represents the line y=x. The rows represent the number of variants included, and the columns indicate the true value of c, the slope between two  vectors used in simulating the data. B, Distribution of type I error (y-axis) of the test results on simulation data under the null hypothesis H0:1=c2 across combinations of phenotypic correlation (x-axis), number of variants (color), and true values of slope c (panels). The dashed horizontal line represents the nominal significance level y=0.05. 28

Figure S9 | ALLSPICE test results on data simulated under the null hypothesis H0:1=c2 across 264 scenarios when 2=0.01 and =0.5. A, QQ plot of test results from the simulated null data described above. The solid black line denotes the line y=x. Rows correspond to the number of variants included, and the columns indicate the true value of the slope parameter c relating the two effect size vectors 1, 2 used in the simulation. B, Distribution of type I error rates (y-axis) under the null hypothesis H0:1=c2 across combinations of phenotypic correlation (x-axis), number of variants (colors), and true values of slope c (panels). The dashed horizontal line represents the nominal significance level y = 0.05. 29

Figure S10 |  ALLSPICE test results with the true value of c on data simulated under the null hypothesis H0:1=c2 across 264 scenarios when 2=1 and =0.8. A, QQ plot of test results from the simulated null data described above. The solid black line denotes the line y=x. Rows correspond to the number of variants included, and the columns indicate the true value of the slope parameter c relating the two effect size vectors 1, 2 used in the simulation. B, Distribution of type I error rates (y-axis) under the null hypothesis H0:1=c2 across combinations of phenotypic correlation (x-axis), number of variants (colors), and true values of slope c (panels). The dashed horizontal line represents the nominal significance level y = 0.05. 30

Figure S11 | ALLSPICE test results on data simulated under the null hypothesis H0:1=c2 across 264 scenarios when variance 2=0.1 and =0.8. A, QQ plot of test results from the simulated null data described above. The solid black line denotes the line y=x. Rows correspond to the number of variants included, and the columns indicate the true value of the slope parameter c relating the two effect size vectors 1, 2 used in the simulation. B, Distribution of type I error rates (y-axis) under the null hypothesis H0:1=c2 across combinations of phenotypic correlation (x-axis), number of variants (colors), and true values of slope c (panels). The dashed horizontal line represents the nominal significance level y = 0.05. 31

Figure S12 | ALLSPICE test results on data simulated under the null hypothesis H0:1=c2 across 264 scenarios when the variance 2=0.01 and the mixture component weight =0.8. A, QQ plot of test results from the simulated null data as described above. The solid black line denotes the line y=x. Rows correspond to the number of variants included, and the columns indicate the true value of the slope parameter c, relating the two effect size vectors 1, 2 used in the simulation. B, Distribution of type I error rates (y-axis) under the null hypothesis H0:1=c2 across combinations of phenotypic correlation (x-axis), number of variants (color), and true values of slope c (panels). The dashed horizontal line represents the nominal significance level y=0.05. 32

Figure S13 | Distribution of empirical statistical power of ALLSPICE test under the alternative hypotheses (2=0.1). Empirical power (y-axis) was evaluated in simulated data across phenotypic correlation (color), number of variants (x-axis), and mixture component weights  (columns), indicating the probability that a variant effect is drawn from the non-zero component, modeled as a standard normal distribution N(0, 2), where 2=0.1. Results are shown for three alternative scenarios: A, independent effect sizes across phenotypes; B, correlated but non-proportional effect sizes; C, non-linear relationship between effect sizes. 34

Figure S14 | Distribution of empirical statistical power of ALLSPICE test under the alternative hypotheses (2=0.01). Empirical power (y-axis) was evaluated in simulated data across phenotypic correlation (color), number of variants (x-axis), and mixture component weights  (columns), indicating the probability that a variant effect is drawn from the non-zero component, modeled as a standard normal distribution N(0, 2), where 2=0.01. Results are shown for three alternative scenarios: A, independent effect sizes across phenotypes; B, correlated but non-proportional effect sizes; C, non-linear relationship between effect sizes. 35

**ALLSPICE testing on Genebass results 36**

Figure S15 | Summary of 11,810 ALLSPICE results across 359 continuous phenotypes on rare variants with AF < 1x10-4. A, QQplot of pALLSPICE filtered phenotypes with Ncases> 300,000, stratified by functional annotation group (colors and columns). B, Relationship between phenotypic correlation (x-axis) and pALLSPICE (negative log-scaled on y-axis). Each point represents a single pair of association; columns indicate the functional annotation group, and point size reflects the number of variants included in each test. The horizontal dashed line in A and B denotes the Bonferroni-corrected significance threshold 4.23 ×10-6. C, Boxplots showing number of variants (log10-scaled y-axis) across three significance categories (x-axis): not significant (pALLSPICE ≥ 0.05), nominally significant (4.23 ×10-6 ≤ pALLSPICE<0.05), or strictly significant (pALLSPICE < 4.23 ×10-6 ), stratified by functional annotation group (colors and columns). 37

Figure S16 | Leave-one-variant-out (LOVO) cross-validation results for rare variants in ALB (AF < 1x10-4). Each panel shows the comparison of variant effect sizes on albumin and calcium levels within ALB, stratified functional annotation group (colors and columns). Each point represents a variant in ALB. The x- and y-axes show the effect sizes of variants for albumin and calcium levels, respectively. Point shapes indicate the nominal significance of single-variant association tests (pGWAS < 0.05). The numbers in the bottom right of each panel report the pALLSPICE for the corresponding annotation group, computed using rare variants with AF < 1x10-4. Point size reflects the influence of each variant in the LOVO analysis: A |log10(cLOVO/cALLvariants )|  and B |log10(PLOVO/PALLvariants )| . In panel A, transparent points indicate variants whose removal increases test significance, whereas solidpoints indicate variants whose removal decreases significance. 39

Figure S17 | Cross-phenotype associations with ALPL. A, Correlation matrix of four phenotypes associated with ALPL across functional groups. B, Distribution of ALPL missense variants associated with alkaline phosphatase levels (pink) or phosphate levels but not alkaline phosphatase levels (green), mapped onto the 3D structure of ALPL protein (PDB ID: 7YIV). C, Distribution of ALPL missense variants affecting alkaline phosphatase and phosphate levels in the same direction (pink) or opposite directions (green), mapped onto the 3D structure of ALPL protein (PDB ID: 7YIV). 41

Figure S18 | Summary of 11,791 ALLSPICE results across 359 continuous phenotypes on ultra-rare variants with AC < 5, restricted to association pairs with phenotypes having phenotypic correlation r < 0.9. A, QQplot of pALLSPICE stratified by functional annotation group (colors and columns). The horizontal dashed line denotes the Bonferroni-corrected significance threshold (p-value=4.24 ×10-6). B, Relationship between phenotypic correlation (x-axis) and pALLSPICE (-log10p, y-axis). Each point represents a single pair; columns indicate functional annotation group, and point size reflects the number of variants included in each test. C, Boxplots showing number of variants (log10-scaled y-axis) across three significance categories (x-axis): not significant (pALLSPICE ≥ 0.05), nominally significant (4.24 ×10-6 ≤ pALLSPICE<0.05), or strictly significant (pALLSPICE < 4.24 ×10-6 ), stratified by functional annotation group (colors and columns). D, Heatmap of pALLSPICE for strictly significant association pairs: with phenotype pairs on the x- and y-axes (genes labeled accordingly). Cell color indicates -log10(pALLSPICE), shown separately by functional annotation group. 43

Figure S19 | Ultra-rare ALB missense variants (AC < 5) on ALB protein structure. A, Distribution of ALB missense variants associated with calcium level but not albumin level (pink), associated with albumin but not calcium level (green), and calcium binding sites (orange) mapped onto the 3D structure of ALB protein (PDB ID: 1AO6). B, Comparison of pairwise distances from missense variants associated exclusively with calcium- (pink) or albumin- (green) levels to calcium binding sites on ALB protein structure in the 3D space. Colors in panel B are defined the same as in panel A. C, Distribution of ALB missense variants associated with either albumin or calcium level that affect the two biomarker levels in the same (blue) or opposite (red) direction(s), together with calcium binding sites (orange) on the 3D structure of ALB protein (PDB ID: 1AO6). D, Comparison of pairwise distances from missense variants to calcium binding sites on the ALB protein structure in the 3D space. Colors in panel D are defined the same as in panel C. 44

Figure S20 | Leave-one-variant-out (LOVO) cross-validation results for ultra-rare variants in ALB (AC < 5). Each panel shows the comparison of variant effect sizes on albumin and calcium levels within ALB, stratified functional annotation group (colors and columns). Each point represents a variant in ALB. The x- and y-axes show the effect sizes of variants for albumin and calcium levels, respectively. Point shapes indicate the nominal significance of single-variant association tests (pGWAS < 0.05). The numbers in the bottom right of each panel report the pALLSPICE for the corresponding annotation group, computed using rare variants with AC < 5. Point size reflects the influence of each variant in the LOVO analysis: A |log10(cLOVO/cALLvariants )| and B |log10(PLOVO/PALLvariants )| . In panel A, transparent points indicate variants whose removal increases test significance, whereas solidpoints indicate variants whose removal decreases significance. 45

References 46

### Permutation and Poisson dispersion test

To assess whether cross-phenotype associations across independent phenotypes occur more frequently than expected by chance, we permuted the $p_{burden}$ for 58,559 high-quality gene-annotation pairs within each of the 239 approximately independent phenotypes, repeating this 100 times within each functional annotation group. For each permutation, we quantified cross-phenotype associations as the proportion of genes with two or more significant associations and compared this to the observed data. Across all four functional annotation groups, the proportion of genes with cross-phenotype associations in the permuted data was substantially lower than observed ($p < 10^{-100}$; mean proportion of permuted genes = 0.07% < 0.5%; **Figure S1A**), indicating an excess of cross-phenotype associations relative to the permutation-based null.

We then repeated this analysis after stratifying genes by combinations of functional annotation groups and cumulative allele frequency (CAF) intervals. Similar enrichment of cross-phenotype associations was observed across most of the sixteen CAF-annotation combinations (**Figure S1B**), suggesting that the excess number of associations is not explained by allele frequency alone. Deviations from this pattern are observed in combinations with insufficient observations. In particular, predicted loss-of-function (pLoF) variants rarely attain high CAF values due to strong selective constraint, leading to sparse observations in higher CAF bins. Conversely, for missense and synonymous groups, their generally higher CAFs reduce the number of genes in the lower CAF bins, leading to less significant differences between the observed and permuted proportions in the corresponding categories.

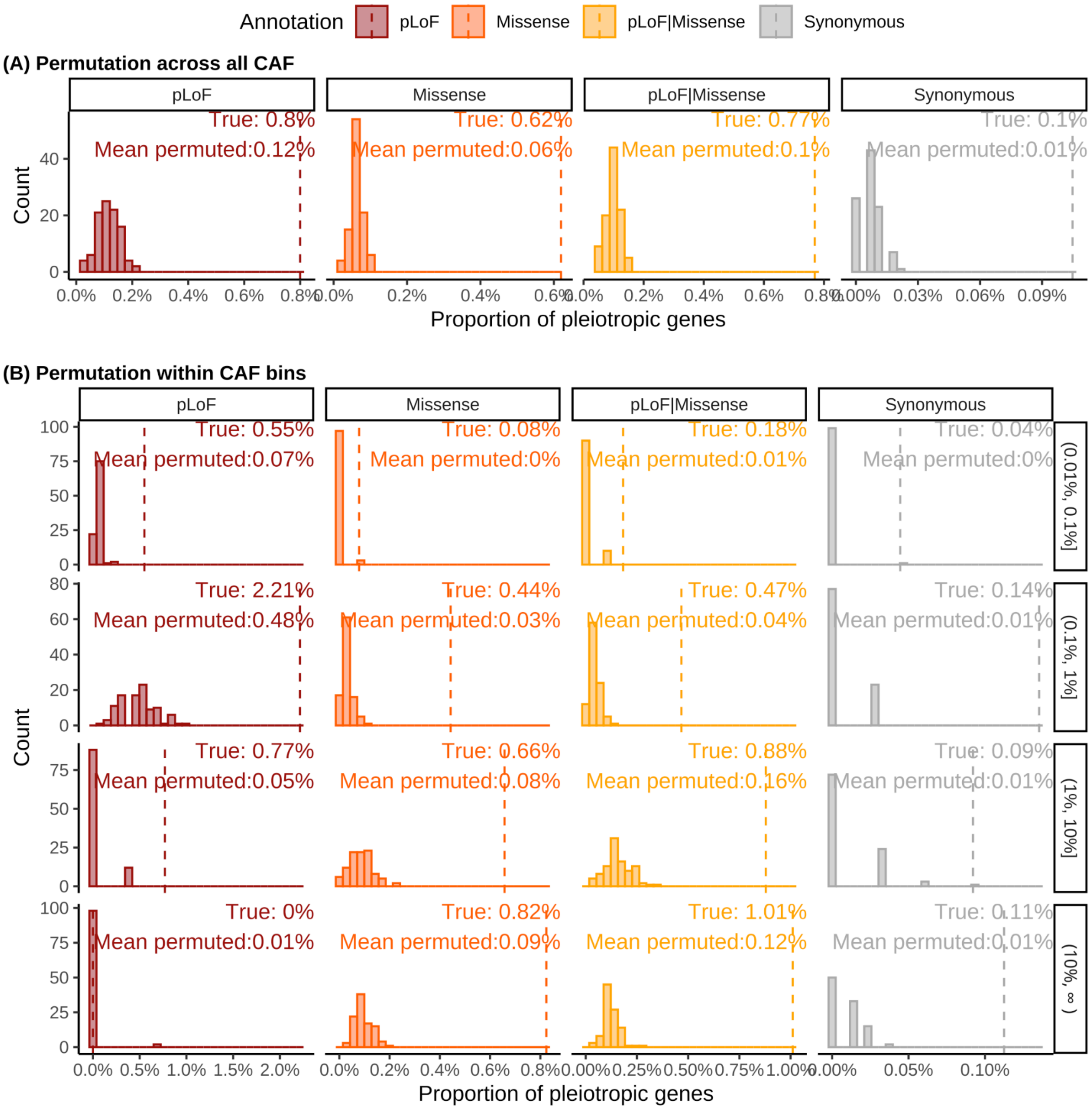


**Figure S1** | Permutation results. **A,** Distribution of the proportion of genes with cross-phenotype associations (x-axis) across permutations for each functional annotation group (columns and colors). **B,** Distribution of the proportion of genes with cross-phenotype associations across permutations, stratified by functional annotation group (columns and colors) and cumulative allele frequency interval (rows). The vertical dashed line in both **A** and **B** denotes the observed proportion in real gene-level association data.

To further evaluate the potential frequency-dependent ascertainment bias, we compared the observed number of associations per gene to expectations under a Poisson model. Specifically, under the null hypothesis, the count of associations for a given functional category within gene $i$ is expected to follow a Poisson distribution with category-specific mean, allowing for heterogeneity across categories defined by functional annotation or CAF. Across functional annotation groups, the observed counts exhibited significant overdispersion relative to the Poisson expectation (Poisson overdispersion test; $p < 10^{-100}$), indicating greater variance than expected given the mean (**Table S1**).

| | **pLoF** | **pLoF + Missense** | **Missense** | **Synonymous** |
|---|---|---|---|---|
| Mean($N_{associations}$) | 0.048 | 0.045 | 0.037 | 0.012 |
| Variance($N_{associations}$) | 0.151 | 0.093 | 0.069 | 0.025 |
| $N_{gene}$ | 7,752 | 16,507 | 17,112 | 17,188 |
| $\chi^2$ Statistic | 24250.00 | 33912.00 | 31830.27 | 34493.39 |
| p-value$_{\chi 2\text{-test}}$ | $< 1x10^{-100}$ | $< 1x10^{-100}$ | $< 1x10^{-100}$ | $< 1x10^{-100}$ |

**Table S1** | Poisson dispersion test for the number of associations per gene-annotation pair. The test evaluates whether the observed distribution of association counts per gene is consistent with a Poisson model under the assumption of independent associations. A significant p-value here indicates overdispersion relative to the Poisson expectation, reflecting an excess of cross-phenotype relative to what would be expected if trait associations were independent. The $\chi^2$ statistic is computed as $\sum_{i=1}^{N_{gene}} \frac{(X_i - \bar{X})^2}{\bar{X}}$ within each functional annotation group, where $\bar{X}$ denotes the mean association count across genes, with the number of associations being $X_i$.

### Pervasive extent of gene-level cross-phenotype associations

Preliminary examination of Genebass rare variant burden test results revealed widespread gene-level associations across multiple traits (**Figure 2A**). Because the detection of cross-phenotype associations of genes is sensitive to statistical power, we further examined how functional annotation and CAF influence the identification of genes with cross-phenotype and trait-specific associations. We stratified genes by four functional annotation groups and four CAF intervals, and then compared the proportion of genes with cross-phenotype associations across these strata.

As previously shown[1], the number of genes with at least one association increases monotonically with CAF, largely reflecting increasing statistical power. However, this trend does not consistently extend to the number of cross-phenotype or trait-specific genes. Genes with associations driven by pLoF variants are more frequently identified at CAF < 1%, whereas a larger number of both cross-phenotype and trait-specific genes are observed for missense variants at higher CAF, particularly above 0.1%. In contrast, synonymous variants show consistently low prevalence of genes with cross-phenotype associations across CAF intervals (**Figure S2**).

Within missense and synonymous groups, the number of cross-phenotype and trait-specific genes increases with CAF, whereas the pLoF group shows the opposite trend across the frequency spectrum (**Figure S2**). Finally, logistic regression analyses assessing the effects of CAF and coding sequence (CDS) length on association status, performed both among genes with at least one association and among all genes, did not identify significant effects for either variable across functional annotation groups at the nominal level ($p < 0.05$). The only exception was a small but statistically significant effect of CDS length on the likelihood of pLoF gene being associated with more than mone phenotypes when considering all genes (p-value = $9.95 \times 10^{-3}$; estimate = $6.26 \times 10^{-5}$) (**Table S2**). These results suggest that gene length has at most a modest influence on association status, with little evidence that CAF alone systematically drives cross-phenotype associations after accounting for association status.

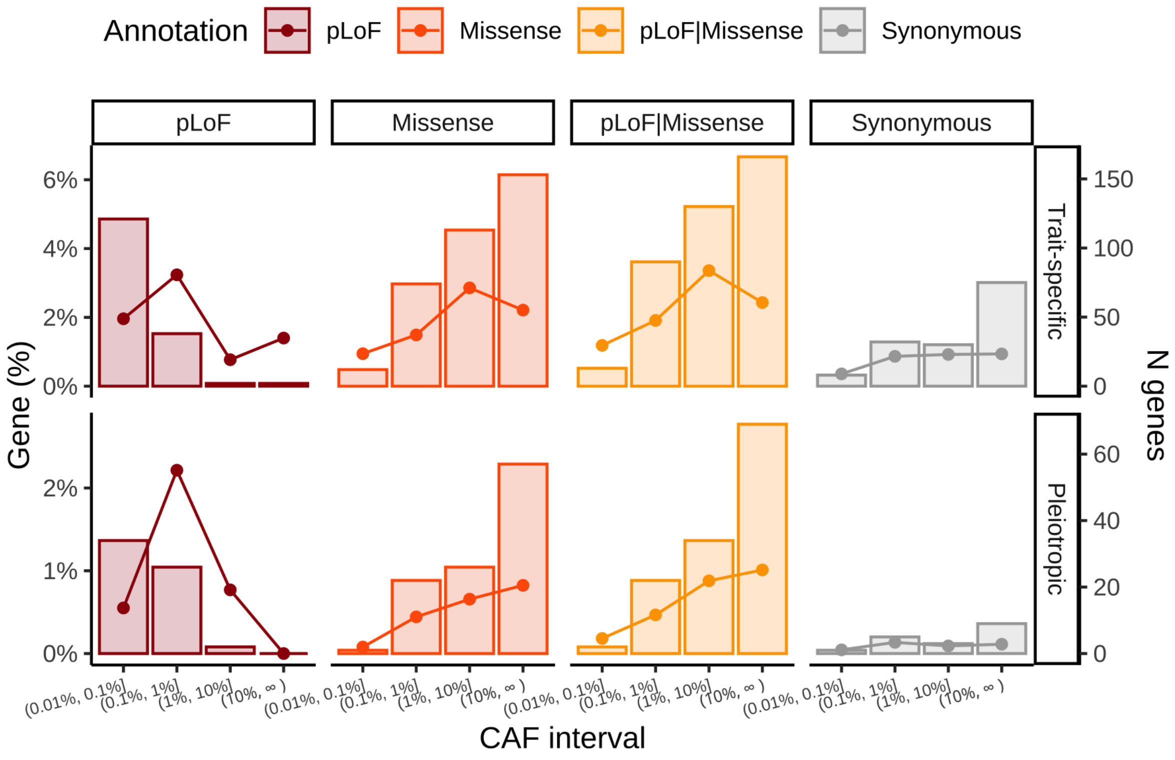


**Figure S2** | Proportion (dot and lines; left y-axis) and count (bars; right y-axis) of genes with single-trait and cross-trait associations (rows) among 239 approximately independent traits ($r^2 < 0.1$), across four CAF intervals (x-axis), stratified by functional annotation group (columns and colors). Proportions are calculated relative to the total number of genes is the total number of genes within each category.

| Annotation | Term | Estimate | std.error | Statistics | P-value |
|---|---|---|---|---|---|
| **pLoF** *($N_{assoc} > 0$)* | (Intercept) | -1.21 | 0.236 | -5.14 | 2.75E-07 |
| | CAF | -2.19 | 9.01 | -0.243 | 8.08E-01 |
| | cds_length | 0.0000834 | 0.0000490 | 1.70 | 8.89E-02 |
| **pLoF + missense** *($N_{assoc} > 0$)* | (Intercept) | -1.14 | 0.153 | -7.48 | 7.52E-14 |
| | CAF | 0.0805 | 0.149 | 0.540 | 5.89E-01 |
| | cds_length | -0.0000051 | 0.0000608 | -0.0834 | 9.34E-01 |
| **Missense** *($N_{assoc} > 0$)* | (Intercept) | -1.17 | 0.163 | -7.21 | 5.75E-13 |
| | CAF | -0.0508 | 0.161 | -0.315 | 7.53E-01 |
| | cds_length | 0.00000965 | 0.0000631 | 0.153 | 8.78E-01 |
| **Synonymous** *($N_{assoc} > 0$)* | (Intercept) | -2.26 | 0.382 | -5.92 | 3.31E-09 |
| | CAF | -0.218 | 0.477 | -0.457 | 6.48E-01 |
| | cds_length | 0.000120 | 0.000121 | 0.989 | 3.23E-01 |
| **pLoF** *($N_{assoc} \geq 0$)* | (Intercept) | -4.90 | 0.146 | -33.7 | 1.04E-248 |
| | CAF | -3.72 | 5.95 | -0.625 | 5.32E-01 |
| | cds_length | 0.0000626 | 0.0000243 | 2.58 | 9.95E-03 |
| **pLoF + missense** *($N_{assoc} \geq 0$)* | (Intercept) | -4.90 | 0.102 | -48.2 | 0 |
| | CAF | 0.0785 | 0.104 | 0.753 | 4.52E-01 |
| | cds_length | 0.0000257 | 0.0000244 | 1.05 | 2.91E-01 |
| **Missense** *($N_{assoc} \geq 0$)* | (Intercept) | -5.12 | 0.110 | -46.4 | 0 |
| | CAF | 0.0825 | 0.115 | 0.720 | 4.71E-01 |
| | cds_length | 0.0000265 | 0.0000256 | 1.03 | 3.01E-01 |
| **Synonymous** *($N_{assoc} \geq 0$)* | (Intercept) | -6.97 | 0.280 | -24.9 | 4.60E-137 |
| | CAF | -0.0536 | 0.394 | -0.136 | 8.92E-01 |
| | cds_length | 0.0000439 | 0.0000487 | 0.902 | 3.67E-01 |

**Table S2** | Logistic regression results assessing the effects of CAF and CDS length on association status. Analyses were performed among genes with at least one association, and among all genes, including those with no associations, stratified by functional annotation group.

To examine whether gene functional importance relates to cross-phenotype effects, we analyzed 37 gene function categories (e.g., essential or haploinsufficient genes;

https://github.com/macarthur-lab/gene_lists). Across annotation groups, pLoF variants show the highest enrichment of genes with cross-phenotype association, followed by missense variants with weaker enrichment, whereas synonymous variants exhibit uniformly low levels of cross-phenotype associations, comparable to background rates. The proportion of genes showing cross-phenotype associations varies across gene functional categories and reflects distinct biological constraints. Olfactory receptor genes, which are generally tolerant to loss-of-function variation and subject to weak selective constraint, are correspondingly depleted for associations and consequently for genes with cross-phenotype associations[3,4]. In contrast, genes with essential biological functions are also depleted for associations, but for a different reason: variants with large, multi-trait effects in these genes are more strongly constrained by purifying selection, reducing their observed associations in population-based studies[5–7]. As a result, essential genes can exhibit lower apparent chance of cross-phenotype assoicaions despite their broad biological roles. Disease-related gene sets, including ACMG genes and BROCA cancer risk panel, show increased chance of cross-phenotype assoicaions for pLoF and missense variants, whereas ClinGen haploinsufficient genes are particularly enriched for cross-phenotype pLoF associations, consistent with dosage sensitivity when a single functional copy of a gene is insufficient (**Figure S3**).

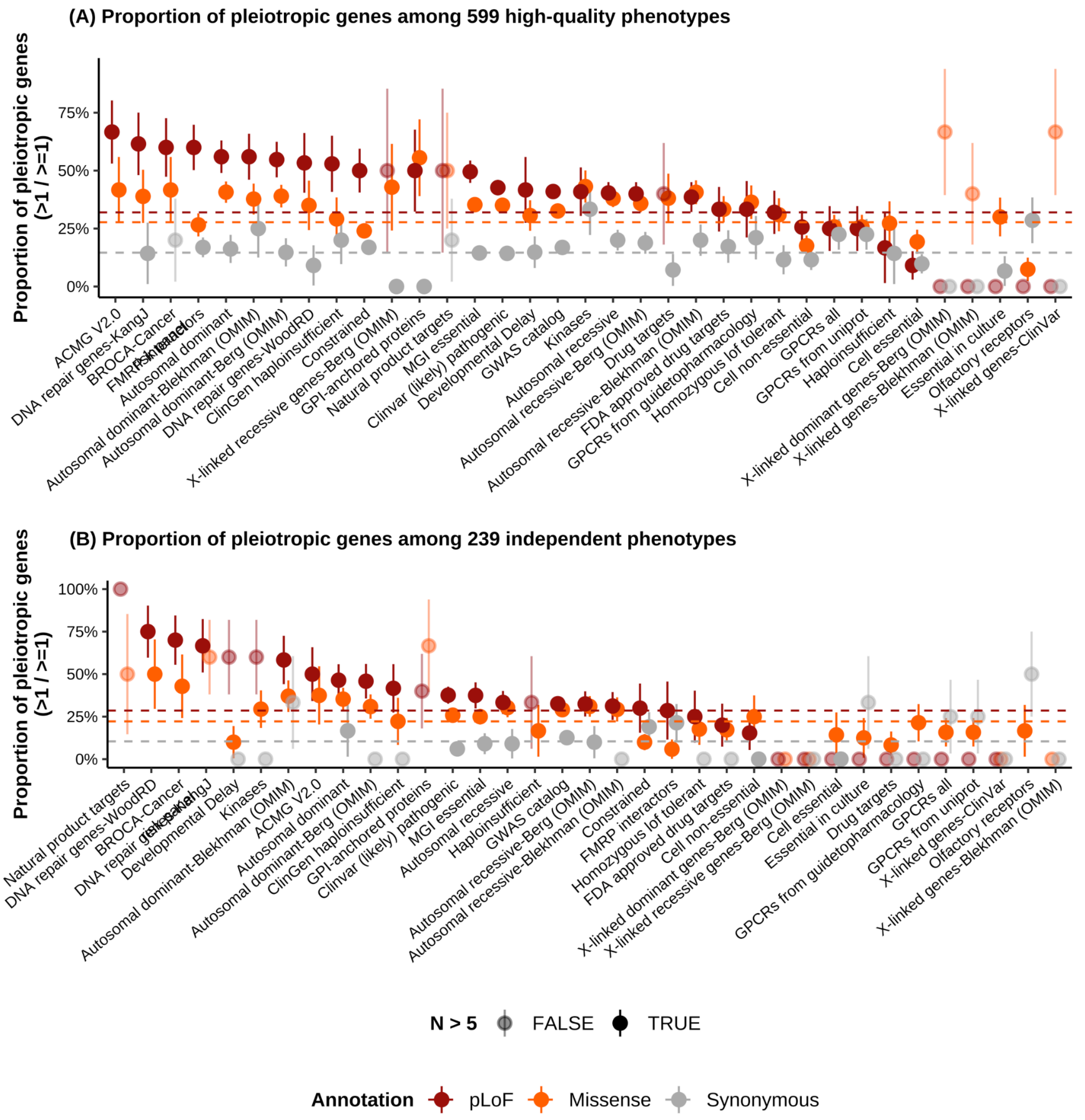


**Figure S3** | Gene-level cross-phenotype associations across all 37 functional gene categories. For each category and functional annotation group, the proportion of genes with cross-phenotype associations is computed as the number of genes with two or more associations divided by the number of genes with at least one association (**SuppTable3**). Transparent points indicate categories with fewer than 5 genes tested. The dashed line denotes the corresponding proportion computed across all genes. Error bars represent standard errors. **A,** Associations across the 599 curated phenotypes; **B,** associations across the 239 approximately independent phenotypes ($r^2 < 0.1$).

To further characterize molecular properties associated with cross-phenotype associations, we queried the STRING database[8] using the R API (STRINGdb, v2.14.3) and computed the mean number of protein-protein interactions (PPIs) for genes with no association, trait-specific association (one association), or cross-phenotype associations (> 1 association), stratified by functional group. Only high-confidence interactions were retained based on the combined STRING score ( > 0.7)[9]. For pLoF and missense variants, genes with cross-phenotype associations show a higher mean number of PPIs compared with genes with zero or one association. In contrast, for synonymous variants, the mean number of PPIs remains similar across association status (**Figure S4**).

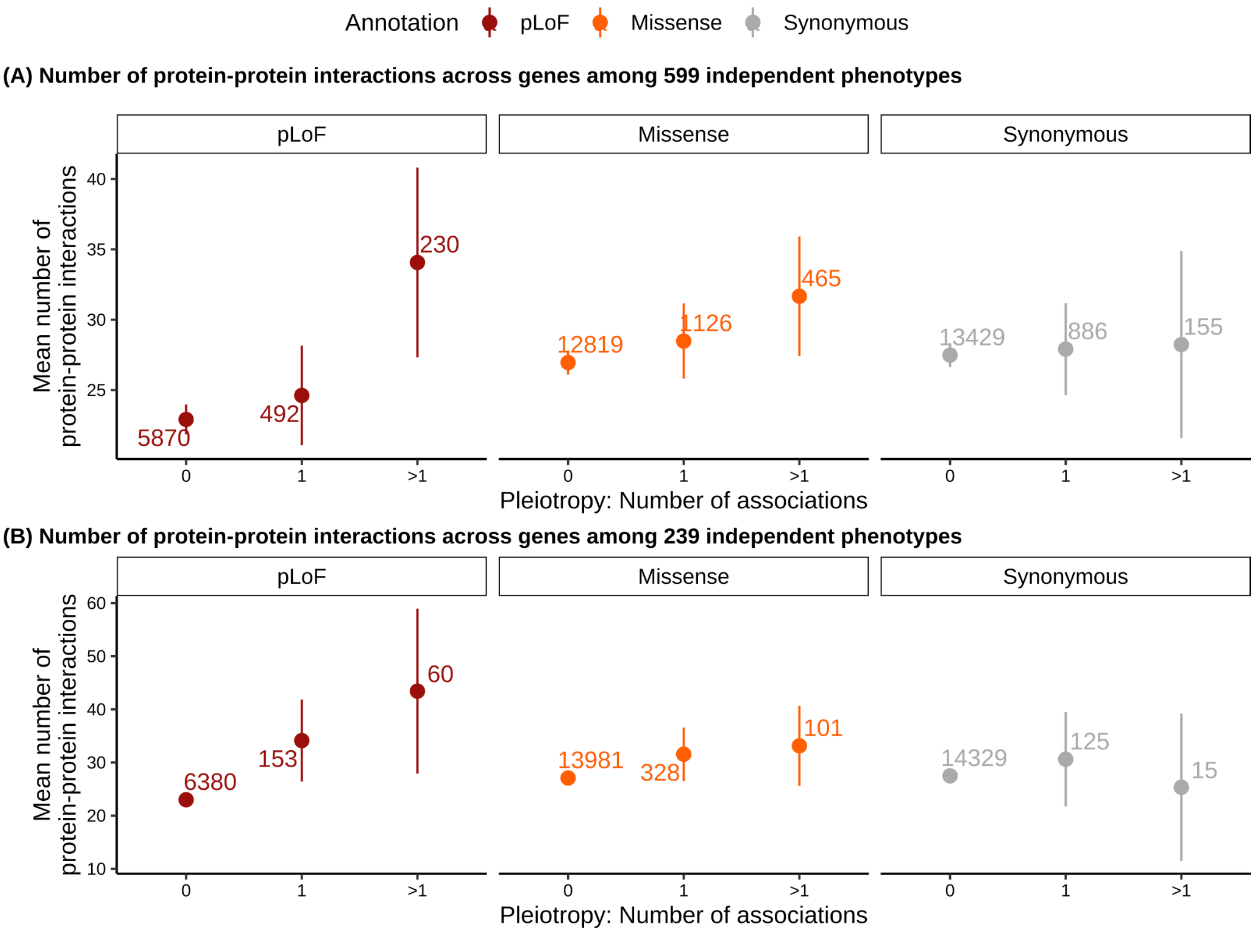

**Figure S4** | Number of protein-protein interactions across association categories. The mean number of protein-protein interactions (y-axis) is shown for genes associated with no association, a single association, or multiple associations (x-axis), stratified by functional annotation group (colors and columns). Numbers above points indicate the total number of genes in each group, and error bars denote 95% confidence intervals. **A,** Associations across the 599 curated phenotypes; **B,** associations across the 239 approximately independent phenotypes ($r^2 < 0.1$).

### Pleiotropy manifests in diverse forms: allelic series and domain-level associations

One form of pleiotropy is an allelic series, in which mutations within the same gene produce a spectrum of related phenotypic outcomes, often reflecting differences in functional severity or molecular mechanisms, such as pLoF versus missense variants[30]. In practice, allelic series are often interpreted within a shared phenotypic pathway, where perturbation of the same gene affects both a proximal continuous trait and a related clinical endpoint, as exemplified by *PCSK9* variants influencing LDL cholesterol levels and coronary heart disease risk[31]. Here, focusing on biomarkers and disease outcomes, we investigated allelic series in gene-based association results by identifying gene-phenotype associations that are significant for either pLoF and missense variants, but not both, highlighting heterogeneity in variant effects across functional annotation groups.

Across all 599 phenotypes (without restricting by phenotypic correlation), we identified 447 genes associated with more than one phenotype for either pLoF or missense group, among which 31 genes show associations with different sets of phenotypes between their pLoF and missense groups (**SuppTable2**). For example, pLoF variants in *ATM*, a well-documented tumor suppressor gene, are associated with six non-hematologic cancer diagnoses and agranulocytosis, a condition that can arise as a side effect of chemotherapy [32,33], whereas missense variants in *ATM* are associated with three erythrocyte-related indices, which may capture cancer-related anemia rather than overt malignancy[34]. Similarly, *PKD1*, *SLC34A3*, and *TMPRSS6* show pLoF associations with diseases but missense associations with blood biomarkers. On the other hand, *FAM234A*, *JAK2*, and *MICA* show the opposite pattern:

missense associations with disease and pLoF associations with blood biomarkers (**Table S3**, **Figure S5** and **SuppTable4**). These differential associations are consistent with the possibility that different variant classes can have different functional consequences within the same gene.

Motivated by the observation that different variant classes within the same gene can be associated with distinct phenotypic sets, we examined the extent to which gene-level associations extend across broader phenotypic domains. Across six phenotypic domains constructed from the 599 curated phenotypes (**Figure S6A**), we identify 191 gene-annotation pairs exhibiting multi-domain associations (**Figure S6C**). Among gene-annotation pairs with at least one association, more deleterious functional classes showed a higher proportion of multi-domain associations (12.2% for pLoF, 7.5% for missense, and 3.4% for synonymous) and a lower proportion of trait-specific associations (50.1% for pLoF, 57.1% for missense, and 69.7% for synonymous) (**Table S5 and Figure S6B**). A similar relationship between functional deleteriousness and association breadth was observed across 17 disease-level domains (**Table S6 and Figure S7**).

Among genes with multi-domain associations, pLoF variants in genes *GIGYF1* and *KDM5B* are each associated with the most (n = 4) distinct phenotypic domains. For *KDM5B*, these associations span atrial fibrillation and flutter[35], several blood biomarkers, physical measures including hand grip strength[36], spirometry traits, and fluid intelligence score[37]. For *GIGYF1*, associations include hand grip strength[36], current tobacco smoking, biomarkers such as HbA1c and glucose, and type 2 diabetes, consistent with prior reports linking *GIGYF1* to endocrine-related traits[38] **Table S4**). As for examples of domain-specific association, missense variants in *OCA2*, a gene involved in pigmentation biology through its role in melanin production[39], are associated with both skin changes related to chronic exposure to non-ionizing radiation ($p_{burden} = 6.15 \times 10^{-8}$) and basal cell carcinoma ($p_{burden} = 2.02 \times 10^{-6}$). In addition, ankylosing spondylitis and iridocyclitis share gene-level associations, including missense variants in *NELFE* ($p_{ankylosing\ spondylitis} = 3.44 \times 10^{-13}$; $p_{iridocyclitis} = 9.01 \times 10^{-14}$), *PSMB9* ($p_{ankylosing\ spondylitis}$

= 1.95 × $10^{-8}$; $p_{iridocyclitis}$ = 5.00 × $10^{-8}$), and *SLC44A4* ($p_{ankylosing\ spondylitis}$ = 1.45 × $10^{-7}$; $p_{iridocyclitis}$ = 3.6 × $10^{-7}$). Among these, *PSMB9* encodes a subunit of the immunoproteasome, which processes intracellular proteins into peptides for presentation by MHC Class I molecules, including HLA-B27, a genetic risk factor for ankylosing spondylitis[40,41]. *NELFE*, which regulates transcriptional elongation, and *SLC44A4*, a transporter gene, are not disease-specific but may reflect shared regulatory or systemic genetic influences contributing to these inflammatory phenotypes.

| Gene | pLoF-only associations | Missense-only associations | Both pLoF and missense associations |
|---|---|---|---|
| *ATM* | •**Cancer diagnosed by doctor**<br>•**C25 Malignant neoplasm of pancreas**<br>•**C50 Malignant neoplasm of breast**<br>•**C77 Secondary and unspecified malignant neoplasm of lymph nodes**<br>•**C78 Secondary malignant neoplasm of respiratory and digestive organs**<br>•**C79 Secondary malignant neoplasm of other sites**<br>•**D70 agranulocytosis** | •Mean reticulocyte volume<br>•Mean corpuscular volume<br>•Mean corpuscular haemoglobin | |
| *PKD1* | •**Hypertension**<br>•Urate<br>•Microalbumin in urine<br>•Haemoglobin concentration<br>•Urea<br>•Cystatin C | •Alkaline phosphatase<br>•Monocyte count | •Creatinine |
| *TMPRSS6* | •**D50 iron deficiency anaemia**<br>•Mean corpuscular haemoglobin concentration<br>•Mean sphered cell volume | •Glycated haemoglobin (HbA1c) | •Haemoglobin concentration<br>•Red blood cell (erythrocyte) distribution width<br>•Haematocrit percentage<br>•Mean corpuscular volume<br>•Mean corpuscular haemoglobin |
| *JAK2* | •Eosinophill count<br>•Eosinophill percentage | •**D75 other diseases of blood and blood-forming organs**<br>•Mean platelet (thrombocyte) volume<br>•Red blood cell (erythrocyte) distribution width | •IGF-1<br>•Platelet count<br>•Platelet crit |
| *SLC34A3* | •**N20 calculus of kidney and ureter** | •Urea | •Cystatin C<br>•Creatinine<br>•Phosphate |
| *FAM234A* | •Glucose | •**Diabetes diagnosed by doctor** | |
| *MICA* | •Lymphocyte count | •**Hypothyroidism/myxoedema**<br>•**E03 other hypothyroidism** | |

**Table S3** | Genes for which pLoF variants are associated with disease phenotypes and missense variants with biomarkers and no diseases, or vice versa. Disease phenotypes are shown in bold.

| Phenotypic domain | *GIGYF1* | | *KDM5B* | |
|---|---|---|---|---|
| | Phenotype | $p_{burden}$ | Phenotype | $p_{burden}$ |
| **Diseases** | E11 Type 2 Diabetes | $1.55 \times 10^{-11}$ | I48 atrial fibrillation and flutter | $2.19 \times 10^{-7}$ |
| | J44 Other Chronic obstructive pulmonary disease | $1.57 \times 10^{-6}$ | | |
| | E03 Other hypothyroidism | $1.20 \times 10^{-7}$ | | |
| **Biomarkers** | Apolipoprotein A | $1.27 \times 10^{-6}$ | IGF-1 | $2.29 \times 10^{-12}$ |
| | Apolipoprotein B | $1.18 \times 10^{-12}$ | Red blood cell (erythrocyte) distribution width | $1.15 \times 10^{-6}$ |
| | Cholesterol | $2.57 \times 10^{-16}$ | Creatinine | $1.34 \times 10^{-6}$ |
| | LDL direct | $6.45 \times 10^{-13}$ | | |
| | Total protein | $2.08 \times 10^{-6}$ | | |
| | HDL cholesterol | $4.69 \times 10^{-8}$ | | |
| | Glycated haemoglobin (HbA1c) | $2.19 \times 10^{-20}$ | | |
| | IGF-1 | $2.03 \times 10^{-6}$ | | |
| | Cytatin C | $6.85 \times 10^{-8}$ | | |
| | Glucose | $1.22 \times 10^{-14}$ | | |
| **Physical measurements** | Peak expiratory flow (PEF) | $7.40 \times 10^{-9}$ | Peak expiratory flow (PEF) | $3.48 \times 10^{-8}$ |
| | Hand grip strength (left) | $2.77 \times 10^{-14}$ | Hand grip strength (left) | $2.15 \times 10^{-12}$ |
| | Hand grip strength (right) | $4.71 \times 10^{-10}$ | Hand grip strength (right) | $7.49 \times 10^{-11}$ |
| | | | Forced expiratory volume in 1 second (FEV1) | $3.37 \times 10^{-10}$ |
| | | | Forced vital capacity (FVC) | $9.81 \times 10^{-8}$ |
| **Mental & cognitive measurements** | | | Fluid intelligence score | $1.38 \times 10^{-6}$ |
| | | | Mean time to correctly identify matches | $1.33 \times 10^{-12}$ |
| **Diet & lifestyle** | Current tobacco smoking | $1.77 \times 10^{-6}$ | | |

**Table S4** | Gene burden associations ($p_{burden} < 2.5 \times 10^{-6}$) of pLoF variants in *GIGYF1* and *KDM5B* across phenotypic domains.

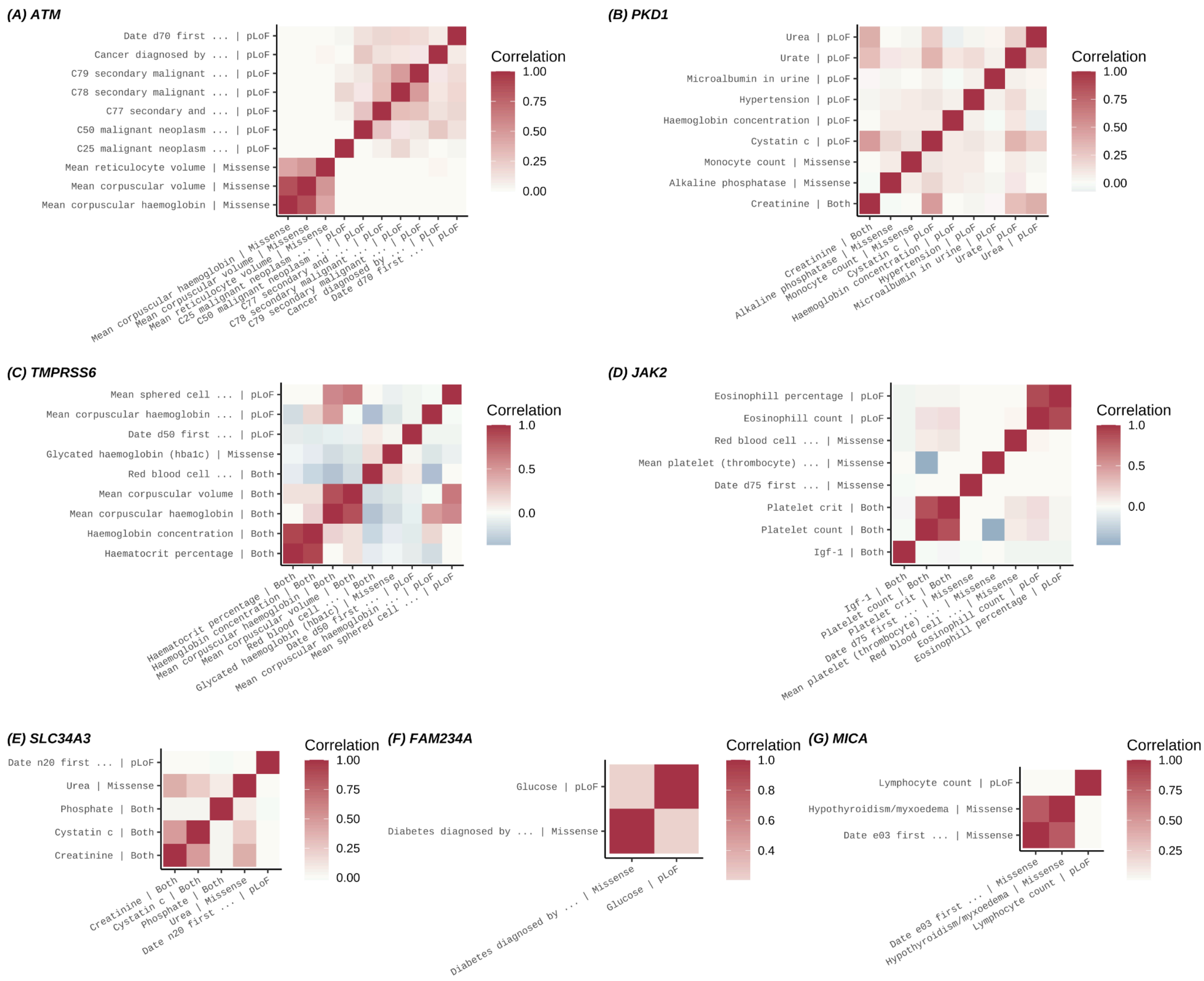


**Figure S5** | Phenotypic correlation matrices for phenotypes listed in **Table 1**. Matrices are shown for phenotypes associated with **A**, *ATM*; **B**, *PKD1*; **C**, *TMPRSS6*; **D**, *JAK2*; **E**, *SLC34A3*; **F**, *FAM234A*; and **G**, *MICA*. Variant annotation groups are indicated alongside the phenotype labels on both the x- and y-axes.

### Genes with cross-domain associations across phenotypic and disease domains

Prior studies have characterized association status at different levels: multi-domain (associated with traits from multiple domains), domain-specific (associated with multiple traits within one domain), and trait-specific (a single trait), and have reported that more than 60% of genes show multi-domain associations through common variants across 27 domains[10]. Based on this framework, more recent work has operationalized phenotype grouping by mapping phenotypes to phecode-based categories, classifying cross-phenotype signals according to whether associated traits fall within a single phecode category or across multiple ones[11]. Following a similar framework, we classified the 599 curated phenotypes into six broad phenotypic domains (**Figure S6A**), and further categorized 234 disease phenotypes into 17 ICD-10-based disease domains (**Figure S7A**). Using these definitions, we identified 191 gene-annotation pairs with cross-domain associations, including 45 pLoF, 77 pLoF+ missense, 58 missense, and 11 synonymous groups (**Figure S6C**). In addition, 38 gene-annotation pairs were associated with multiple disease domains, comprising 12 pLoF, 17 pLoF + missense, 8 missense, and 1 synonymous group (**Figure S7C**).

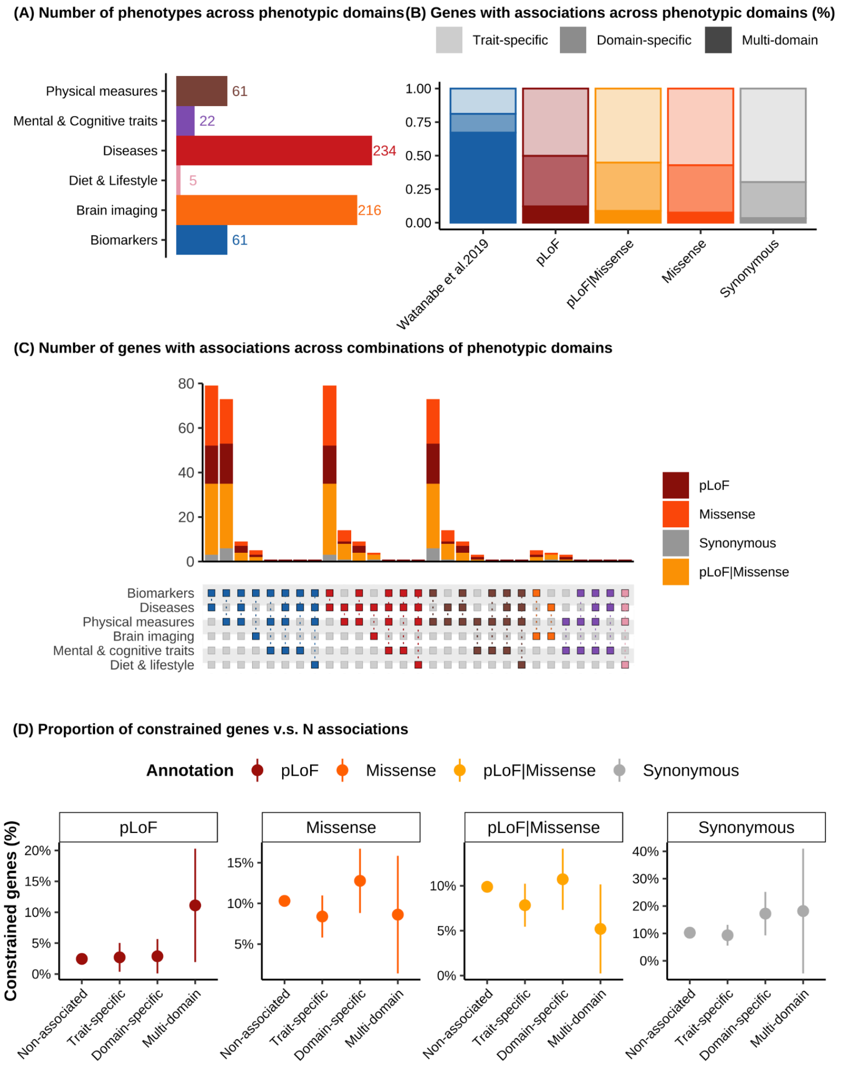


**Figure S6** | Pervasive gene-level cross-domain associations across six broader phenotypic domains: **A,** Number of phenotypes in each of the six phenotypic domains (color-coded); **B,** Proportion of genes with trait-specific, domain-specific, and multi-domain associations defined at the phenotypic domain level, among gene-annotation pairs with at least one association across functional annotation groups (color-coded). The blue bar shows the common variant results from Watanabe et al. 2019[1]; **C,** Number of genes associated with each combination of phenotypic domains. The bars indicate the number of genes with cross-domain associations, as shown in the bottom matrix, where each column represents a domain combination that shares at least one gene-level association. Colors are consistent between panel **A** and the bottom matrix in panel **C**. **D,** Proportion of constrained genes, defined as those in the lowest LOEUF decile (y-axis), with no association, trait-specific, domain-specific, or multi-domain associations (x-axis) across functional annotation groups (color and column). Error bars represent 95% confidence intervals.

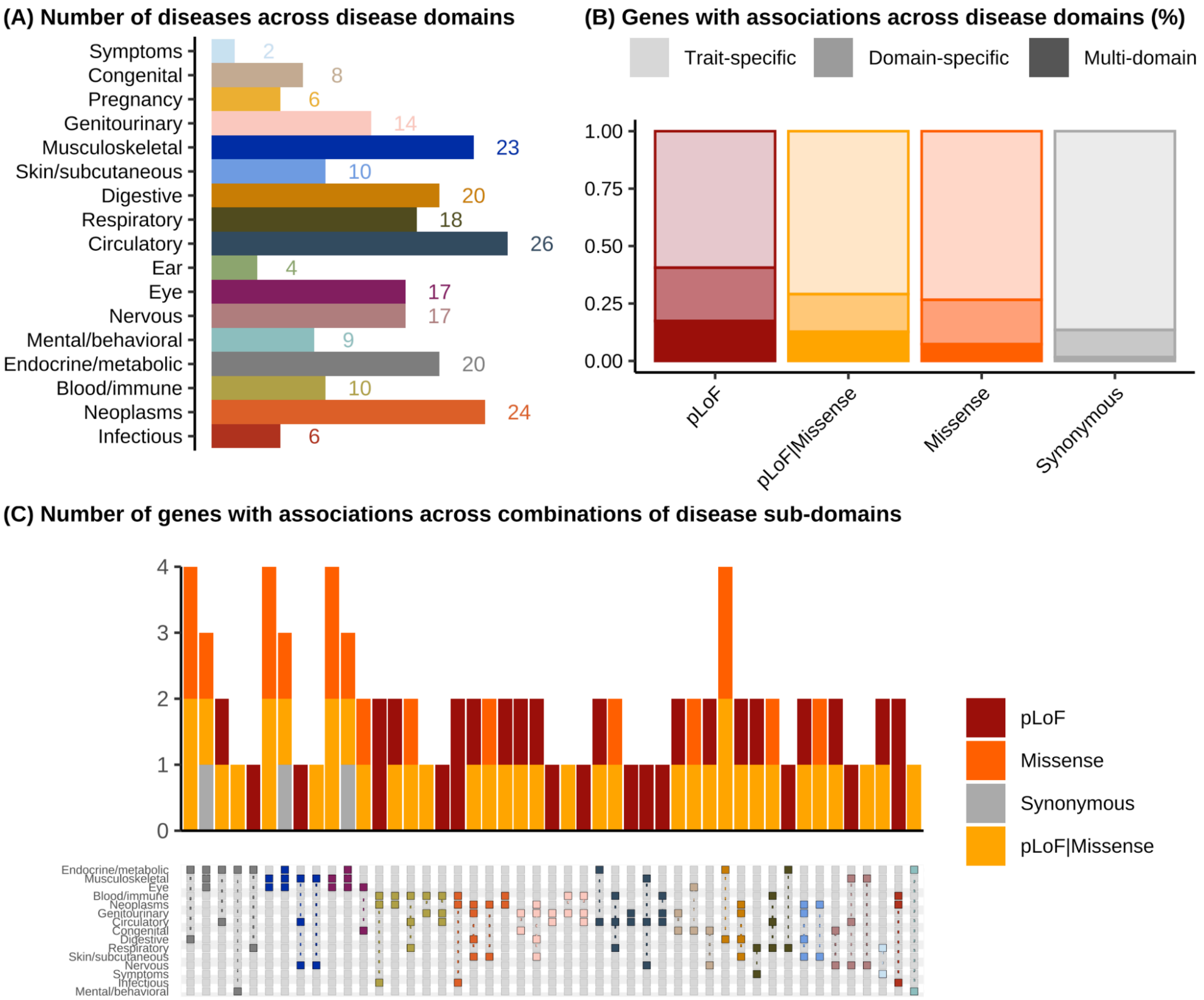


**Figure S7** | Gene-level cross-domain associations across 17 disease sub-domains. **A,** Number of diseases in each of the 17 disease sub-domains; **B,** Proportion of genes with trait-specific, domain-specific, and multi-domain associations at the disease sub-domain level, among gene-annotation pairs with at least one association across, stratified by functional annotation group. **C,** Number of genes associated with each combination of disease sub-domains. Each column of the bottom matrix represents a distinct sub-domain combination sharing gene-level associations. Bars indicate the number of genes with cross-disease-domain associations defined by the corresponding dot combinations in the matrix below. Colors are consistent between panel **A** and the matrix in panel **C**.

We performed a rough comparison between common and rare variants using summary statistics from Wanatabe et al.[10] and Genebass[1]. In Watanabe et al. (2019), 65.9% of genes are associated with at least one trait through common variants, and among these, 81.2% show evidence of cross-phenotype associations. These genes with cross-phenotype associations are further classified into multi-domain (67.17%), domain-specific (14.03%), or trait-specific (18.80%) across 27 phenotypic domains. In contrast, we consider a smaller set of six phenotypic domains from Genebass, and a majority of gene-annotation groups, 95.24% of pLoF, 95.48% of missense, and 98.12% of synonymous, show no associations. Among the associated genes, pLoF variants are more enriched for genes with cross-phenotype associations (49.86%), particularly for multi-domain (12.20%) and domain-specific (37.67%) genes, compared with missense and synonymous variants, while exhibiting the lowest proportion of trait-specific associations (50.14%) (**Figure S6B**; **Table S5**). A similar pattern is observed when association status is evaluated across the 17 disease domains (**Figure S7**; **Table S6**). These results highlight potential differences in the distribution and detection of cross-phenotype associations between common and rare variants. However, direct comparison is limited by substantial differences in study design, phenotype definitions, domain resolution, statistical power, and potential confounding, making pleiotropic metrics from the two studies not directly comparable.

| | Watanabe et al. | | Genebass | | | | | |
|---|---|---|---|---|---|---|---|---|
| | Genes | | pLoF | | Missense | | Synonymous | |
| | N | % | N | % | N | % | N | % |
| **Total in the genome** | 17,518 | 100.00 | 7,752 | 100.00% | 17,112 | 100.00% | 17,188 | 100.00% |
| **Associated** | 11,544 | 65.90 | 369 | 4.76% | 773 | 4.52% | 323 | 1.88% |
| Cross-phenotype | 9,374 | 81.20 | 184 | 49.86% | 332 | 42.95% | 98 | 30.34% |
| Multi-domain | 7,754 | 67.17 | 45 | 12.20% | 58 | 7.50% | 11 | 3.41% |
| Domain-specific | 1,620 | 14.03 | 139 | 37.67% | 274 | 35.45% | 87 | 26.93% |
| Trait-specific | 2,170 | 18.80 | 185 | 50.14% | 441 | 57.05% | 225 | 69.66% |
| **Non-associated** | 5,974 | 34.10 | 7,383 | 95.24% | 16,339 | 95.48% | 16,865 | 98.12% |

**Table S5** | Comparison with common variant results from Watanabe et al. (2019). Shown are the proportion and number of genes with cross-phenotype, multi-domain, domain-specific, and trait-specific associations across six broad phenotypic domains (599 phenotypes), stratified by functional annotation group.

| | Genebass | | | | | |
|---|---|---|---|---|---|---|
| | pLoF | | Missense | | Synonymous | |
| | N | % | N | % | N | % |
| **Total in the genome** | 7,752 | 100.00% | 17,112 | 100.00% | 17,188 | 100.00% |
| **Disease-associated** | 69 | 0.89% | 109 | 0.64% | 59 | 0.34% |
| Cross-phenotype | 28 | 40.58% | 29 | 26.61% | 8 | 13.56% |
| Multi-domain | 12 | 17.39% | 8 | 7.34% | 1 | 1.69% |
| Domain-specific | 16 | 23.19% | 21 | 19.27% | 7 | 11.86% |
| Disease-specific | 41 | 59.42% | 80 | 73.39% | 51 | 86.44% |

**Table S6** | Proportion of genes with cross-phenotype, multi-domain, domain-specific, and trait-specific associations across 17 disease sub-domains, stratified by functional annotation group.

Among multi-domain associations, the biomarker domain shows higher overlap with both disease ($r = 0.179$, $p < 10^{-100}$) and physical measurement ($r = 0.085$, $p < 10^{-100}$) domains. For the other pairs of phenotypic domains, correlations of the number of associations per gene are below 0.05 (**Table S7**).

| | Biomarkers | Brain Imaging | Diet & Lifestyle | Diseases | Mental & Cognitive | Physical measures |
|---|---|---|---|---|---|---|
| Biomarkers | 1 | 0.0140 | 0.0286 | **0.1787** | 0.0151 | **0.0852** |
| Brain Imaging | 0.0013 | 1 | -0.0002 | 0.0042 | -0.00054 | -0.0012 |
| Diet & Lifestyle | 0.0115 | -0.0004 | 1 | 0.0407 | -0.0002 | 0.0078 |
| Diseases | **0.1089** | 0.0121 | 0.0254 | 1 | 0.0083 | 0.0320 |
| Mental & Cognitive | 0.0079 | -0.0014 | -0.0002 | 0.0144 | 1 | 0.0256 |
| Physical measures | **0.0929** | -0.0045 | 0.0237 | 0.0568 | 0.0364 | 1 |

**Table S7** | Correlations of the number of associations per gene across phenotypic domains, the values in the upper triangle are correlations between the number of associations per gene within each pair of domains, the values in the lower triangle are correlations between the binary vector, whether there is any association per gene within each pair of domains.

To assess whether gene-level constraint is associated with domain-level association status, we fitted logistic regression models including gene intolerance to loss-of-function variation (LOEUF) score[39] and CDS length, comparing genes with multi-domain, domain-specific, or trait-specific associations to genes with no associations. For pLoF groups, constrained genes with lower LOEUF values were more likely to exhibit multi-domain (estimate = -1.07; OR $\approx$ 2.94; p-value = $5.67 \times 10^{-3}$). Similar but weaker effects were observed for domain-specific (estimate = -0.517; OR $\approx$ 1.67; p-value = $1.81 \times 10^{-2}$) and trait-specific genes (estimate = -0.518; OR $\approx$ 1.67; p-value = $7.13 \times 10^{-3}$) (**Table S6**). In contrast, CDS length showed only a small positive effect across all categories (multi-domain: OR = 1.000045, p-value = $2.18 \times 10^{-2}$; domain-specific: OR = 1.000054, p-value = $1.33 \times 10^{-2}$; trait-specific: OR = 1.000031, p-value = $7.62 \times 10^{-2}$), indicating a statistically detectable but modest contribution.

| Y | $N_{genes}$ | Term | Estimate | Std.error | Statistics | p-value |
|---|---|---|---|---|---|---|
| **Multi-domain vs. No association** | 45 (7,383) | Intercept | -4.19 | 0.380 | -11.0 | $2.65 \times 10^{-28}$ |
| | | oe_lof_upper | -1.07 | 0.388 | -2.77 | $5.67 \times 10^{-3}$ |
| | | cds_length | 0.000054 | 0.000022 | 2.48 | $1.33 \times 10^{-2}$ |
| **Domain-specific vs. No association** | 139 (7,383) | Intercept | -3.55 | 0.242 | -14.7 | $1.06 \times 10^{-48}$ |
| | | oe_lof_upper | -0.517 | 0.219 | -2.36 | $1.81 \times 10^{-2}$ |
| | | cds_length | 0.000045 | 0.00020 | 2.29 | $2.18 \times 10^{-2}$ |
| **Trait-specific vs. No association** | 185 (7,383) | Intercept | -3.25 | 0.213 | -15.3 | $7.34 \times 10^{-53}$ |
| | | oe_lof_upper | -0.518 | 0.192 | -2.69 | $7.13 \times 10^{-3}$ |
| | | cds_length | 0.000031 | 0.000017 | 1.81 | $7.62 \times 10^{-2}$ |

**Table S8** | Logistic regression results for the impact of constraint level and gene CDS length on the pLoF group of a gene having multi-domain, domain-specific, and trait-specific associations compared to having no association.

ALLSPICE testing on simulated data

We simulated data under the null hypothesis with a fixed sample size of $n$ =1000 and the parameter configurations described below (Methods), performing 100 replicate simulations per parameter combination:

- Number of variants within a gene $m \in \{5,\ 10,\ 20, 100\}$,
- Slope between variant-level effect sizes on two phenotypes $c \in \{0,\ 0.2,\ 0.4,\ 0.6,\ 0.8,\ 1\}$
- Phenotypic correlation between the two phenotypes $r \in \{-1,\ -0.8,\ -0.5,\ -0.2,\ -0.1,\ 0,\ 0.1,\ 0.2,\ 0.5,\ 0.8,\ 1\}$
- Variance of the component normal distribution of effect sizes $\sigma^2 \in \{0.01,\ 0.1,\ 1\}$
- Weight for the mixture distribution of variant effects $\pi \in \{0.5,\ 0.8\}$

We applied ALLSPICE to simulated data across all parameter combinations (**Extended Data Figure 1-2**; **Figure S8-Figure S14; SuppTable 9-20**). Across scenarios with $\sigma^2 = 1$, the test shows a well-calibrated p-value distribution (**Extended Data Figure 1A; Figure S10A**), and a reasonable empirical type I error rate. Although type I error increases slightly with the number of variants, it remains well controlled below 0.1 (**Extended Data Figure 1B; Figure S10B**). When $\sigma^2$ decreases ($\sigma^2 = 0.01\ \&\ 0.1$), test statistics exhibit a mildly deflated pattern relative to $\sigma^2 = 1$, consistent with the smaller scale of effect sizes in these simulations (**Figure S8, Figure S9, Figure S11, Figure S12**). Test results for $\pi = 0.8$ are similar to those for $\pi = 0.5$, indicating limited sensitivity of test calibration to this parameter within the examined range (**Figure S10**, **Figure S11**, **Figure S12**).

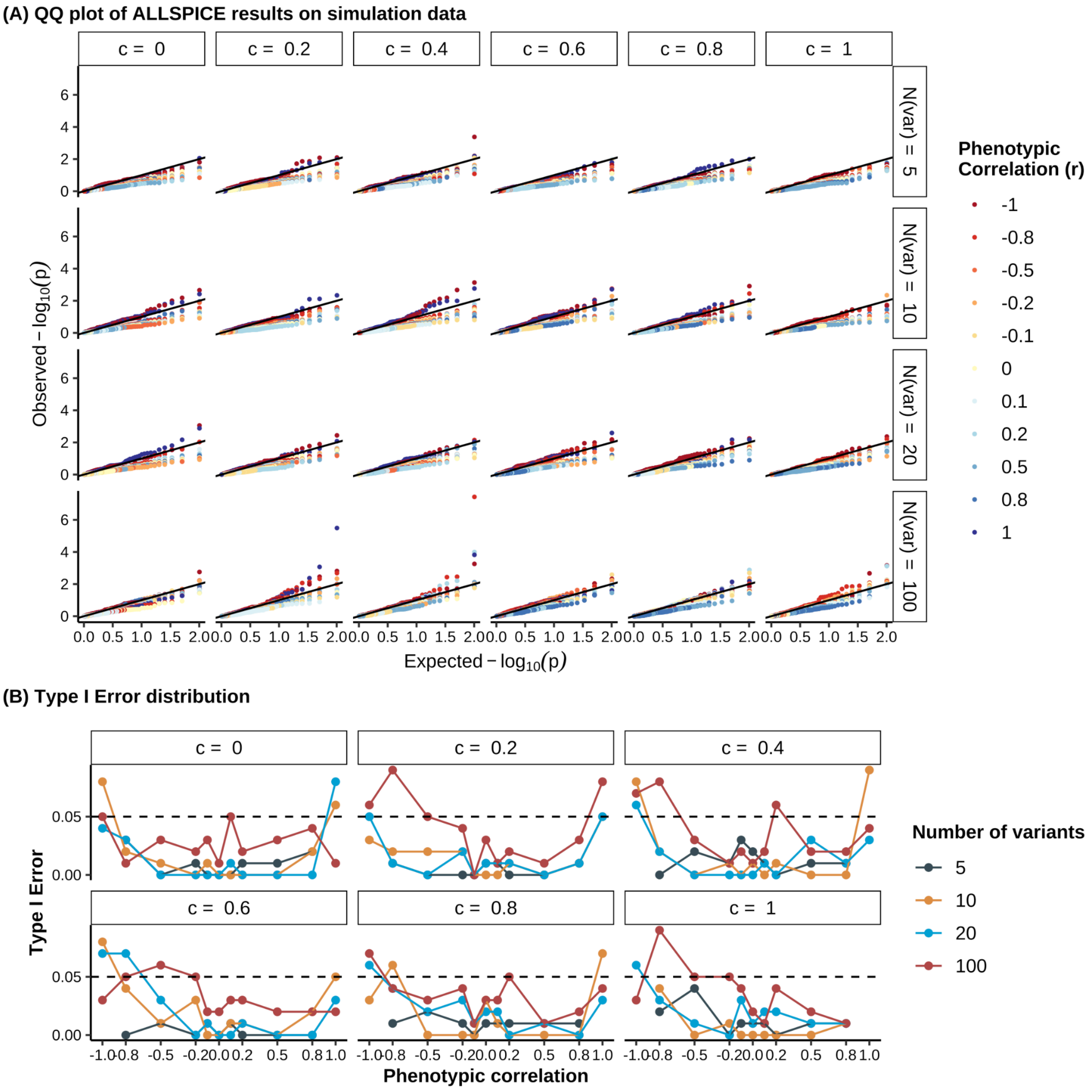


**Figure S8** | ALLSPICE test results on data simulated under the null hypothesis $H_0: \beta_1 = c\beta_2$ across 264 scenarios when $\sigma^2 = 0.1$ and $\pi = 0.5$. **A,** QQ-plot of test results on the simulated data as described above. The solid black line represents the line y=x. The rows represent the number of variants included, and the columns indicate the true value of $c$, the slope between two $\beta$ vectors used in simulating the data. **B,** Distribution of type I error (y-axis) of the test results on simulation data under the null hypothesis $H_0: \beta_1 = c\beta_2$ across combinations of phenotypic correlation (x-axis), number of variants (color), and true values of slope $c$ (panels). The dashed horizontal line represents the nominal significance level y=0.05.

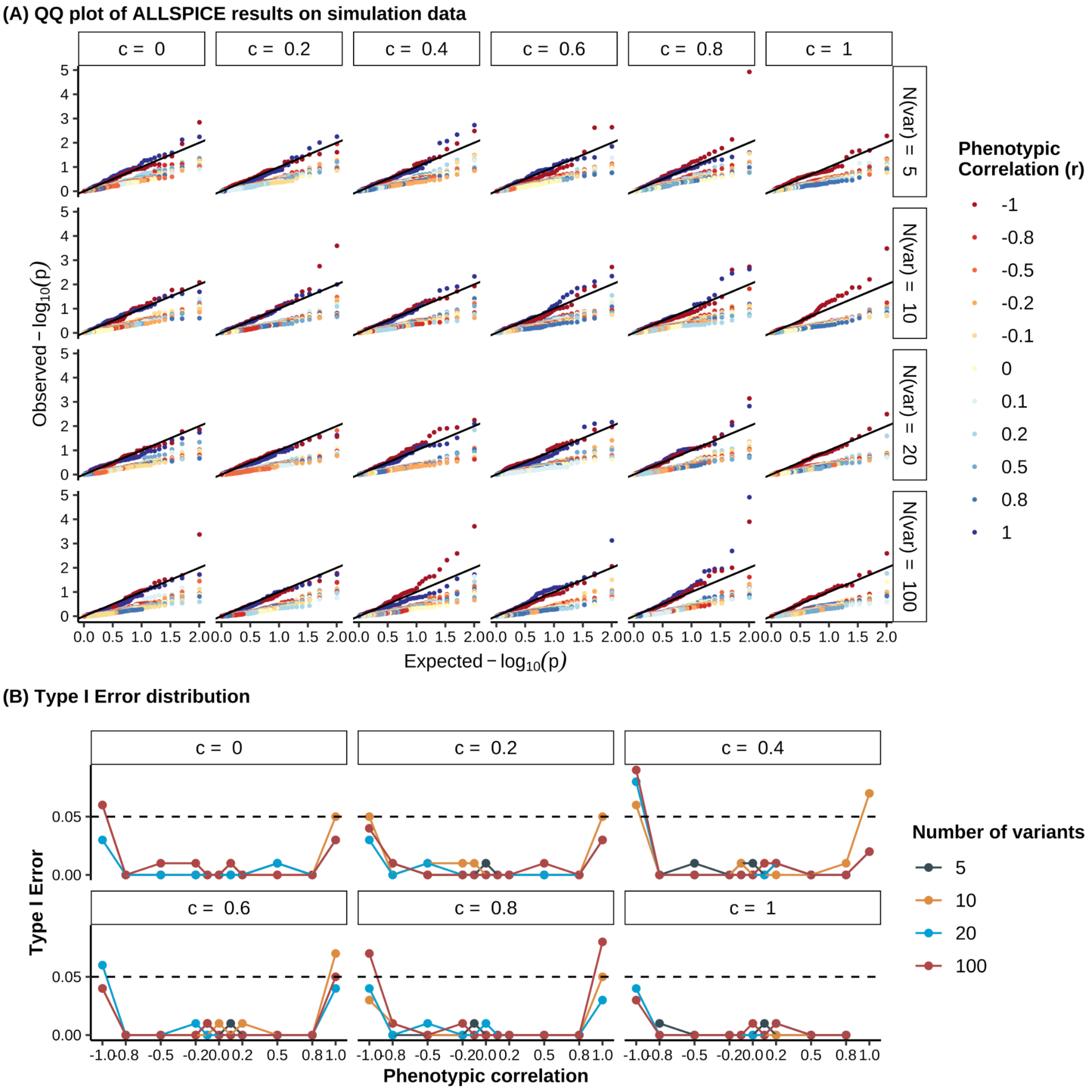


**Figure S9** | ALLSPICE test results on data simulated under the null hypothesis $H_0: \beta_1 = c\beta_2$ across 264 scenarios when $\sigma^2 = 0.01$ and $\pi = 0.5$. **A,** QQ plot of test results from the simulated null data described above. The solid black line denotes the line y=x. Rows correspond to the number of variants included, and the columns indicate the true value of the slope parameter $c$ relating the two effect size vectors $\beta_1$, $\beta_2$ used in the simulation. **B,** Distribution of type I error rates (y-axis) under the null hypothesis $H_0: \beta_1 = c\beta_2$ across combinations of phenotypic correlation (x-axis), number of variants (colors), and true values of slope $c$ (panels). The dashed horizontal line represents the nominal significance level y = 0.05.

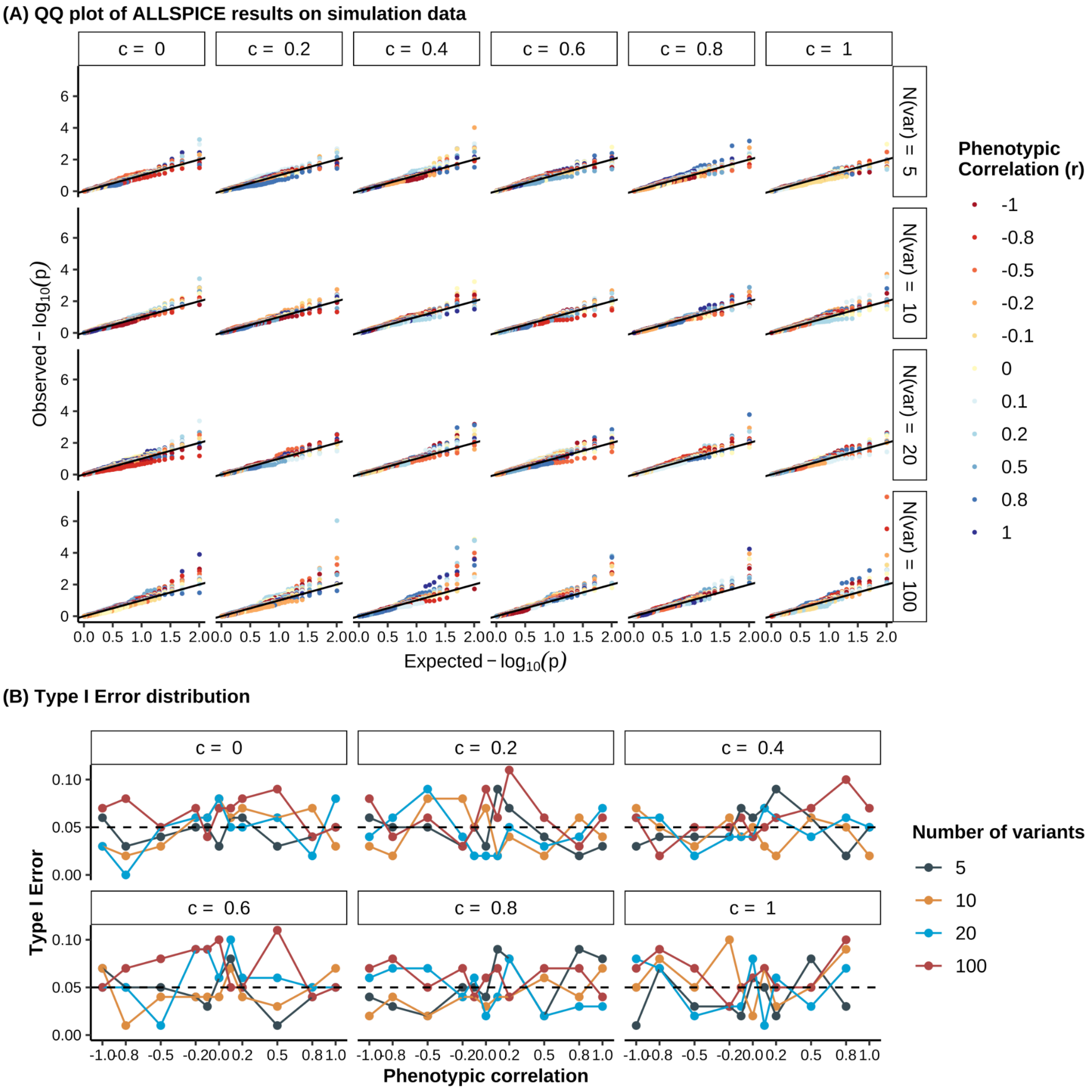


**Figure S10 |** ALLSPICE test results with the true value of $c$ on data simulated under the null hypothesis $H_0: \beta_1 = c\beta_2$ across 264 scenarios when $\sigma^2 = 1$ and $\pi = 0.8$. **A,** QQ plot of test results from the simulated null data described above. The solid black line denotes the line y=x. Rows correspond to the number of variants included, and the columns indicate the true value of the slope parameter $c$ relating the two effect size vectors $\beta_1$, $\beta_2$ used in the simulation. **B,** Distribution of type I error rates (y-axis) under the null hypothesis $H_0: \beta_1 = c\beta_2$ across combinations of phenotypic correlation (x-axis), number of variants (colors), and true values of slope $c$ (panels). The dashed horizontal line represents the nominal significance level y = 0.05.

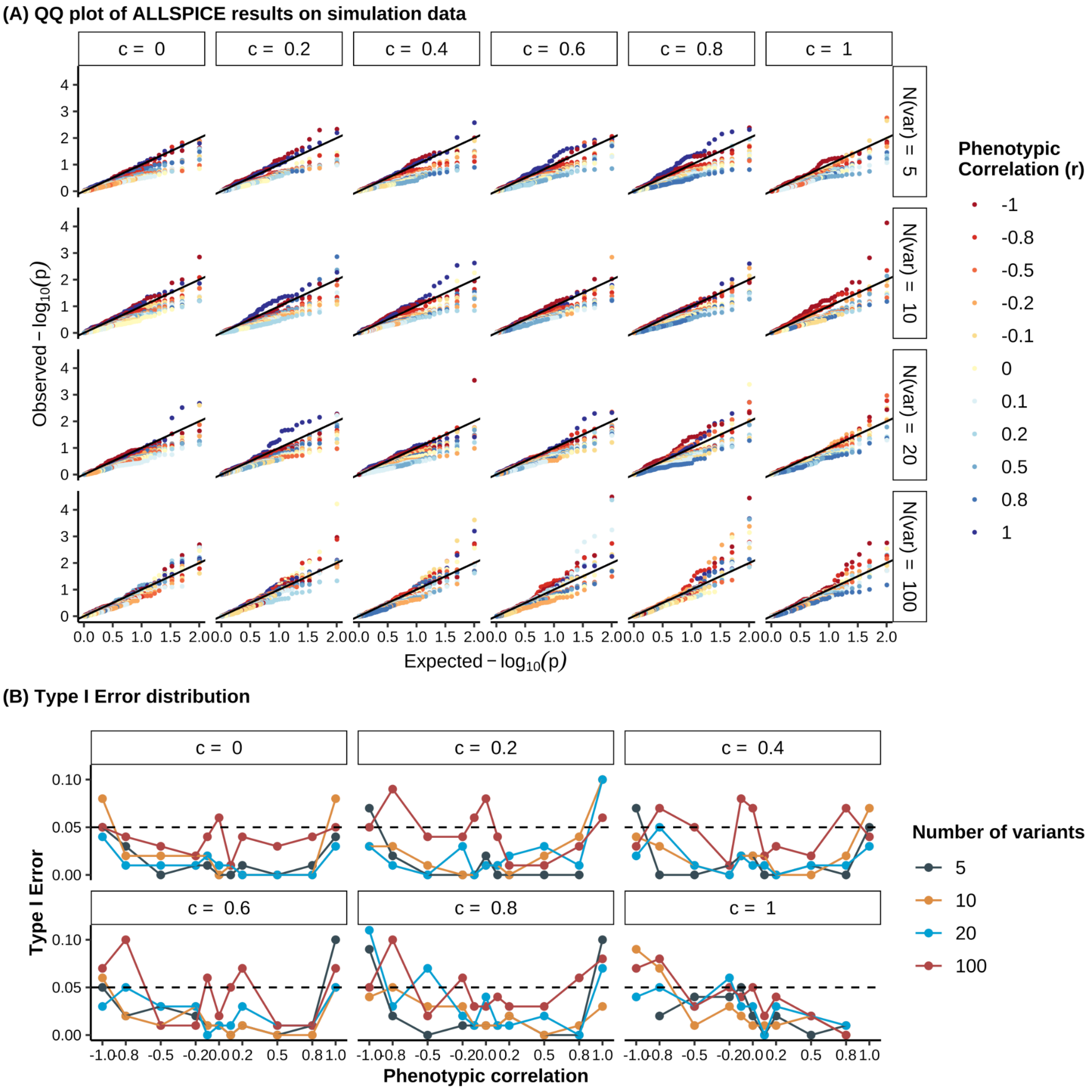


**Figure S11** | ALLSPICE test results on data simulated under the null hypothesis $H_0$: $\beta_1 = c\beta_2$ across 264 scenarios when variance $\sigma^2 = 0.1$ and $\pi = 0.8$. **A,** QQ plot of test results from the simulated null data described above. The solid black line denotes the line y=x. Rows correspond to the number of variants included, and the columns indicate the true value of the slope parameter $c$ relating the two effect size vectors $\beta_1$, $\beta_2$ used in the simulation. **B,** Distribution of type I error rates (y-axis) under the null hypothesis $H_0$: $\beta_1 = c\beta_2$ across combinations of phenotypic correlation (x-axis), number of variants (colors), and true values of slope $c$ (panels). The dashed horizontal line represents the nominal significance level y = 0.05.

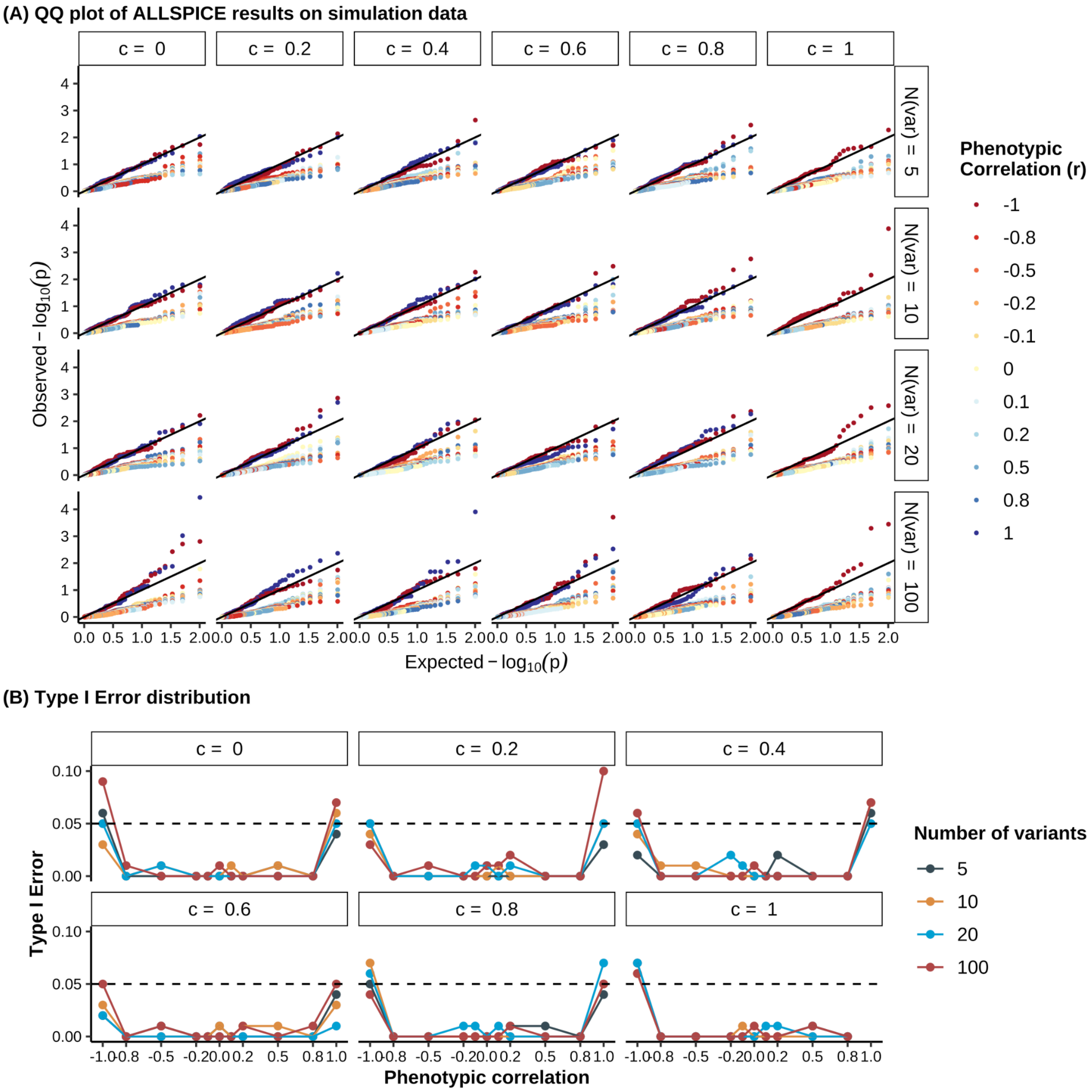


**Figure S12** | ALLSPICE test results on data simulated under the null hypothesis $H_0: \beta_1 = c\beta_2$ across 264 scenarios when the variance $\sigma^2 = 0.01$ and the mixture component weight $\pi = 0.8$. **A,** QQ plot of test results from the simulated null data as described above. The solid black line denotes the line y=x. Rows correspond to the number of variants included, and the columns indicate the true value of the slope parameter $c$, relating the two effect size vectors $\beta_1$, $\beta_2$ used in the simulation. **B,** Distribution of type I error rates (y-axis) under the null hypothesis $H_0: \beta_1 = c\beta_2$ across combinations of phenotypic correlation (x-axis), number of variants (color), and true values of slope $c$ (panels). The dashed horizontal line represents the nominal significance level y=0.05.

We further evaluated empirical statistical power using simulations under the alternative hypothesis (**Methods**), using the same parameter configurations as in the null simulations. For each scenario, we performed 100 replicate simulations and then estimated power as the proportion of significant tests. Power increases with the number of variants, exceeding 0.9 when at least 10 variants are included (**Extended Data Figure 2**). When the variance parameter is reduced ($\sigma^2 = 0.1$ or $0.01$), corresponding to less variability in variant effects, the overall magnitude of simulated effect sizes ($\beta_1$ and $\beta_2$) decreases, leading to a substantial reduction in power to detect significant association pairs (**Figure S13** and **Figure S14**), particularly for scenarios with independent effect sizes.

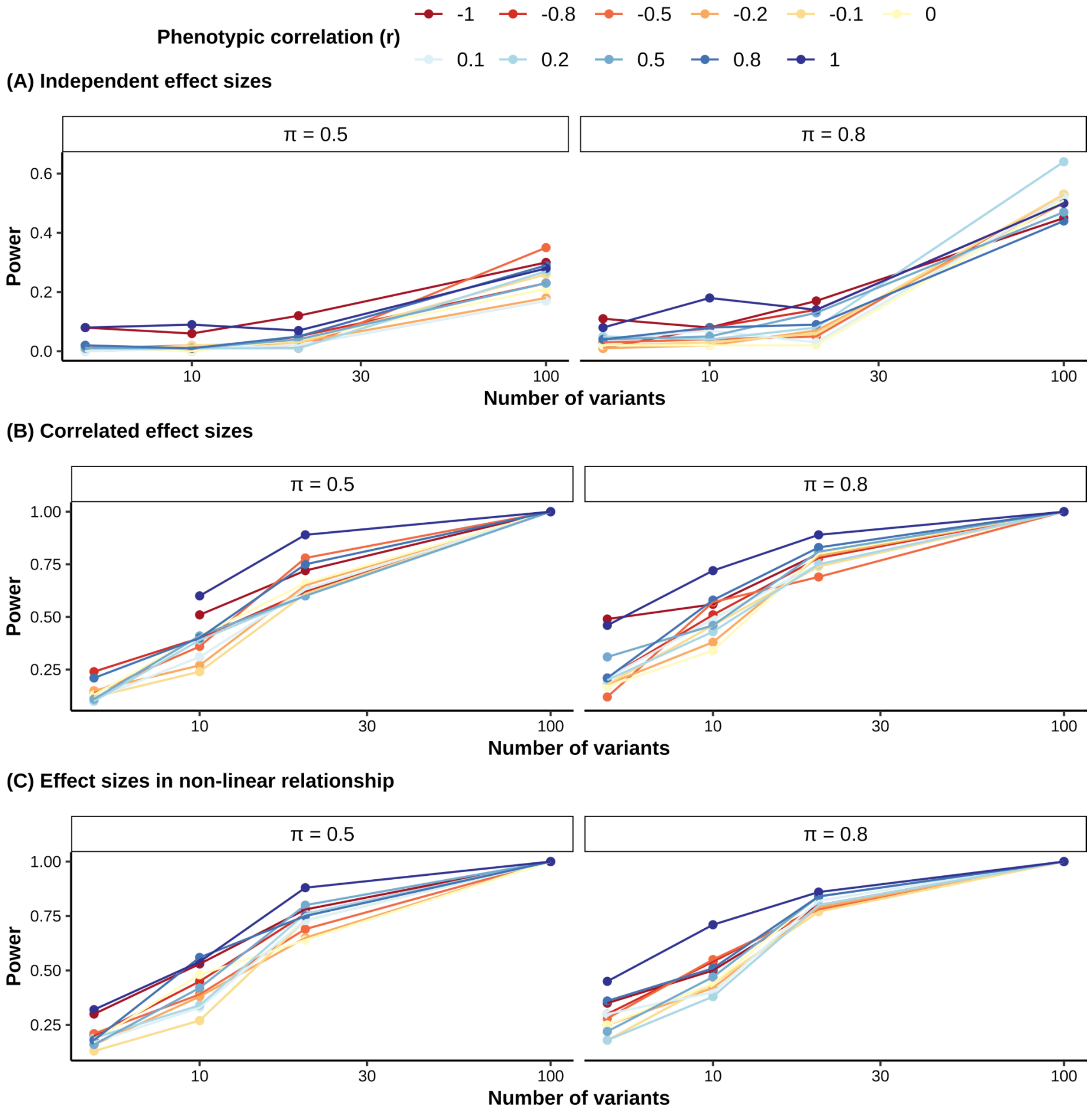


**Figure S13** | Distribution of empirical statistical power of ALLSPICE test under the alternative hypotheses ($\sigma^2 = 0.1$). Empirical power (y-axis) was evaluated in simulated data across phenotypic correlation (color), number of variants (x-axis), and mixture component weights $\pi$ (columns), indicating the probability that a variant effect is drawn from the non-zero component, modeled as a standard normal distribution $N(0, \sigma^2)$, where $\sigma^2 = 0.1$. Results are shown for three alternative scenarios: **A,** independent effect sizes across phenotypes; **B**, correlated but non-proportional effect sizes; **C**, non-linear relationship between effect sizes.

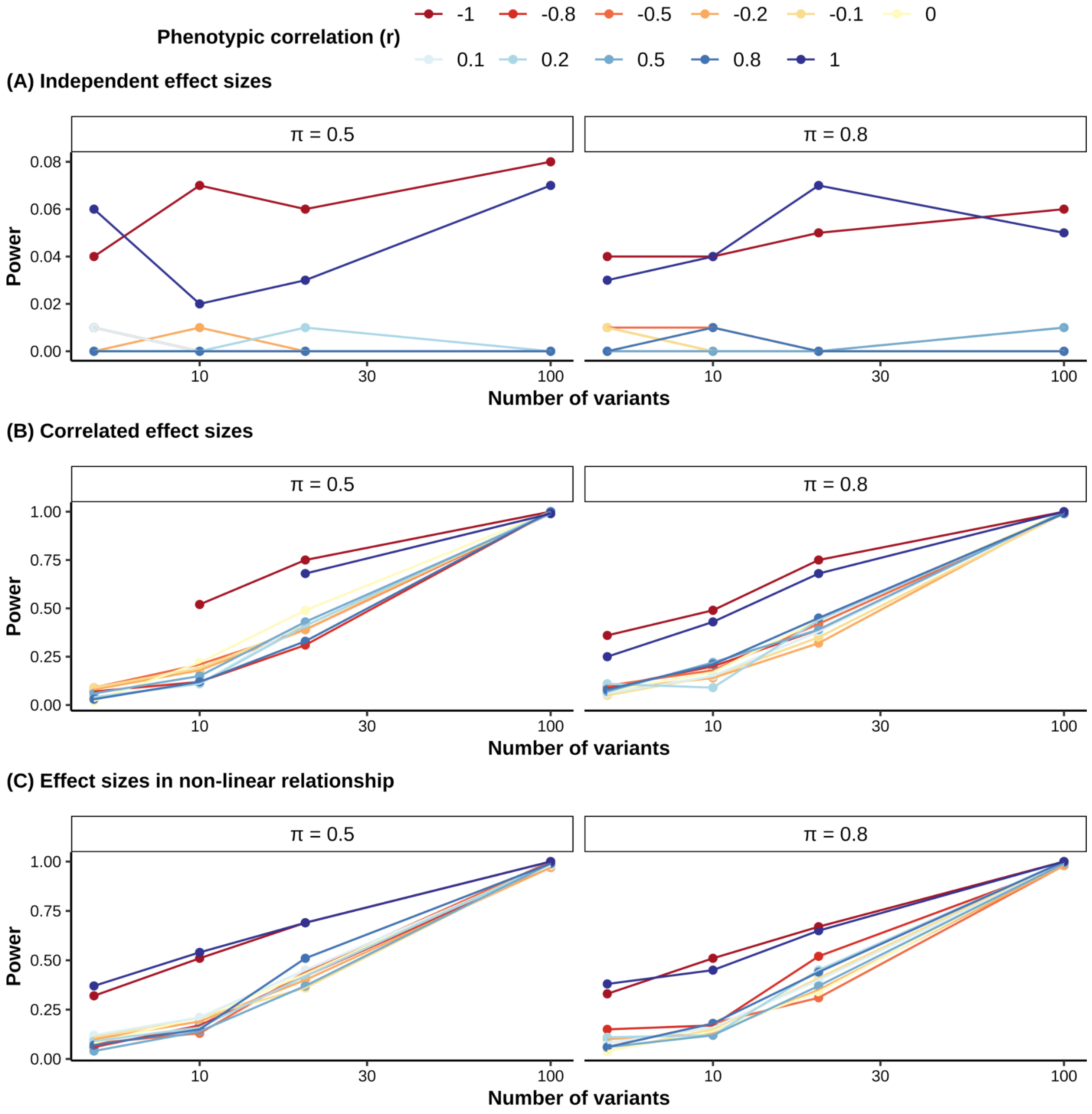


**Figure S14** | Distribution of empirical statistical power of ALLSPICE test under the alternative hypotheses ($\sigma^2 = 0.01$). Empirical power (y-axis) was evaluated in simulated data across phenotypic correlation (color), number of variants (x-axis), and mixture component weights $\pi$ (columns), indicating the probability that a variant effect is drawn from the non-zero component, modeled as a standard normal distribution $N(0, \sigma^2)$, where $\sigma^2 = 0.01$. Results are shown for three alternative scenarios: **A,** independent effect sizes across phenotypes; **B**, correlated but non-proportional effect sizes; **C**, non-linear relationship between effect sizes.

### ALLSPICE testing on Genebass results

In total, we applied ALLSPICE to 11,810 pairs of associations, in which both traits were from 359 continuous traits among the 599 curated phenotypes, and were significantly associated with the corresponding gene-annotation group (**SuppTable 5**). Across rare variants (AF <0.0001), missense variants show a higher proportion of heterogeneous effects across the full range of phenotypic correlations compared with the other functional annotation groups (pLoF 6.2%, missense 11.5%, synonymous 8.4%), consistent with the greater functional heterogeneity of missense variants (**Figure S15A**). Significant association pairs are more frequently observed when the two phenotypes are weakly correlated $(|r| < 0.1)$ or strongly positively correlated $(r > 0.8)$, indicating that phenotypic correlation alone does not fully capture the shared genetic architecture between traits (**Figure S15B**).

ALLSPICE is designed to detect heterogeneous variant effects inconsistent with direct mediation between phenotyes, where the two phenotypes are expected to involve divergent biological pathways. When phenotypic correlation approaches one, variant effect estimates on the two phenotypes become nearly identical, leading to model instability and thus inflation of test statistics (Methods equation 10; **Figure S15B**). Therefore, we restrict downstream analyses to pairs of associations with phenotypic correlation below 0.9 to improve model stability and interpretability.

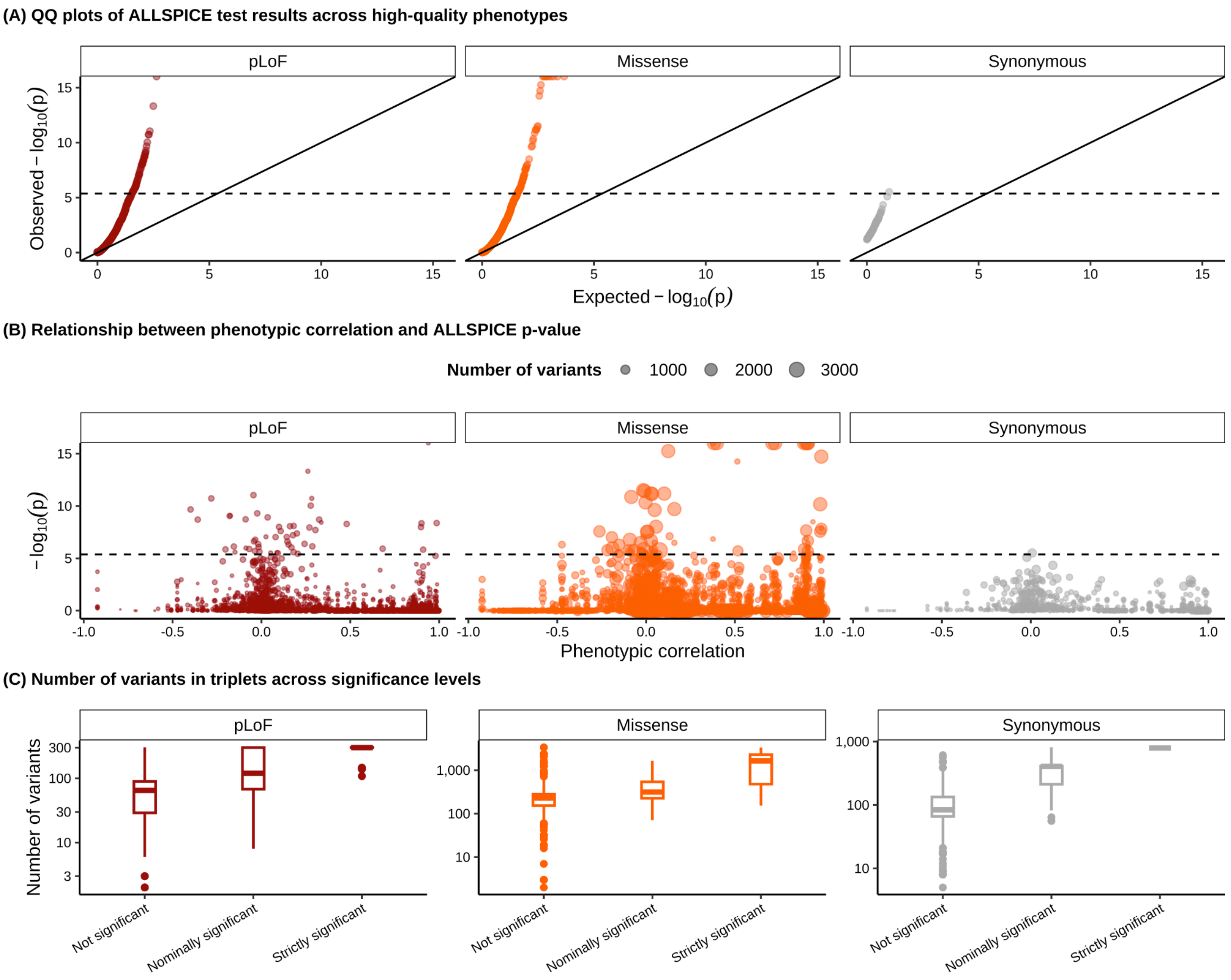


**Figure S15** | Summary of 11,810 ALLSPICE results across 359 continuous phenotypes on rare variants with AF < $1\text{x}10^{-4}$. **A,** QQplot of $p_{ALLSPICE}$ filtered phenotypes with $N_{cases}$> 300,000, stratified by functional annotation group (colors and columns). **B,** Relationship between phenotypic correlation (x-axis) and $p_{ALLSPICE}$ (negative log-scaled on y-axis). Each point represents a single pair of association; columns indicate the functional annotation group, and point size reflects the number of variants included in each test. The horizontal dashed line in **A** and **B** denotes the Bonferroni-corrected significance threshold $4.23 \times 10^{-6}$. **C**, Boxplots showing number of variants ($\log_{10}$-scaled y-axis) across three significance categories (x-axis): not significant ($p_{ALLSPICE} \geq 0.05$), nominally significant ($4.23 \times 10^{-6} \leq p_{ALLSPICE} < 0.05$), or strictly significant ($p_{ALLSPICE} < 4.23 \times 10^{-6}$), stratified by functional annotation group (colors and columns).

To further characterize the sensitivity of ALLSPICE results to individual variants, we performed a leave-one-variant-out (LOVO) analysis for the ALB-albumin-calcium association pair among rare *ALB* variants (AF < $1\text{x}10^{-4}$). ALLSPICE was repeated with one variant removed at a time, and the resulting $p_{ALLSPICE}$ and $c_{MLE}$ were compared with the results obtained using all variants. We found that removing individual variants produces heterogeneous changes in the ALLSPICE test statistics, but did not identify a single variant whose removal alone eliminated the signal (**Figure S16**; **SuppTable 7**). In contrast, when restricting to ultra-rare *ALB* variants (AC < 5), ALLSPICE results became more sensitive to variant inclusion. Removal of 130 variants, largely exhibiting null effect on both phenotypes, increased test significance, whereas removing any of the remaining 83 variants attenuated the test significance, consistent with stronger heterogeneous effects for these variants (**Figure S20**). Notably, exclusion of 26 variants was sufficient to make the test non-significant (**SuppTable 8**), reflecting the increased influence of individual variants when fewer ultra-rare variants are included.

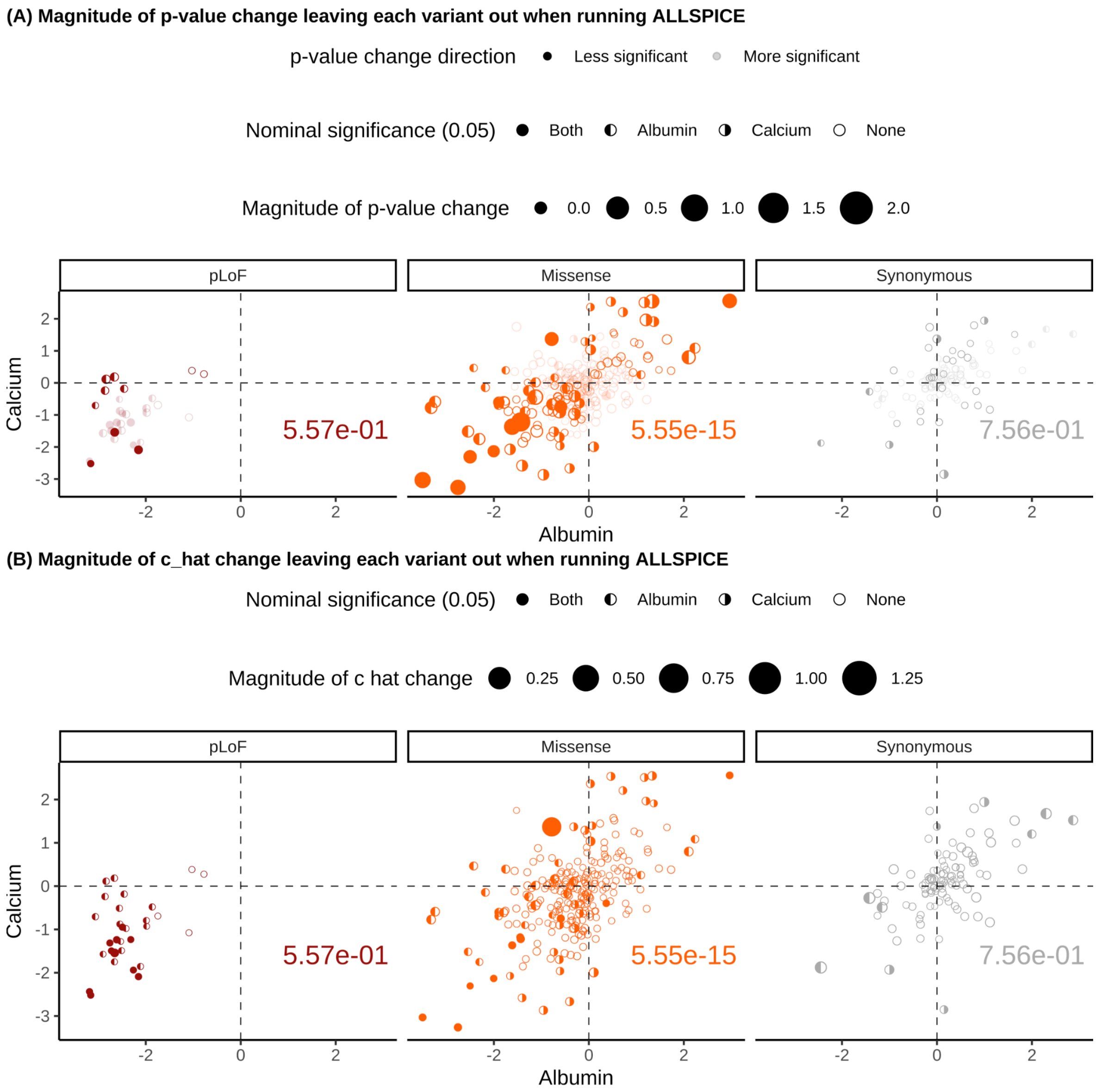


**Figure S16** | Leave-one-variant-out (LOVO) cross-validation results for rare variants in *ALB* (AF < $1\times10^{-4}$). Each panel shows the comparison of variant effect sizes on albumin and calcium levels within *ALB*, stratified functional annotation group (colors and columns). Each point represents a variant in *ALB*. The x- and y-axes show the effect sizes of variants for albumin and calcium levels, respectively. Point shapes indicate the nominal significance of single-variant association tests ($p_{GWAS} < 0.05$). The numbers in the bottom right of each panel report the $p_{ALLSPICE}$ for the corresponding annotation group, computed using rare variants with AF < $1\times10^{-4}$. Point size reflects the influence of each variant in the LOVO analysis: **A** $|log_{10}(c_{LOVO}/c_{ALLvariants})|$ and **B** $|log_{10}(P_{LOVO}/P_{ALLvariants})|$ **.** In panel **A**, transparent points indicate variants whose removal increases test significance, whereas solidpoints indicate variants whose removal decreases significance.

ALLSPICE highlights heterogeneous effects among missense variants in *ALPL* at an allele frequency (AF) threshold of AF < $1x10^{-4}$, motivating further exploration. We investigated the associations of *ALPL* across multiple phenotypes, focusing on alkaline phosphatase and phosphate levels, both of which are significantly associated with pLoF and missense variants in *ALPL*. In addition, missense variants in *ALPL* show significant associations with C-reactive protein and calcium levels. Phenotypic correlations among these four traits vary in magnitude and, in some cases, direction, indicating heterogeneous cross trait effect of *ALPL* (**Figure S17A**). Missense variants were mapped onto the 3D structure of ALPL (PDB ID: 7YIV) to assess their spatial distribution relative to effects on alkaline phosphatase and phosphate levels. These variants do not show a clear spatial clustering within the protein structure, suggesting that their phenotypic associations are not confined to specific structural regions, but rather dispersed across different structural domains (**Figure S17B** and **C**).

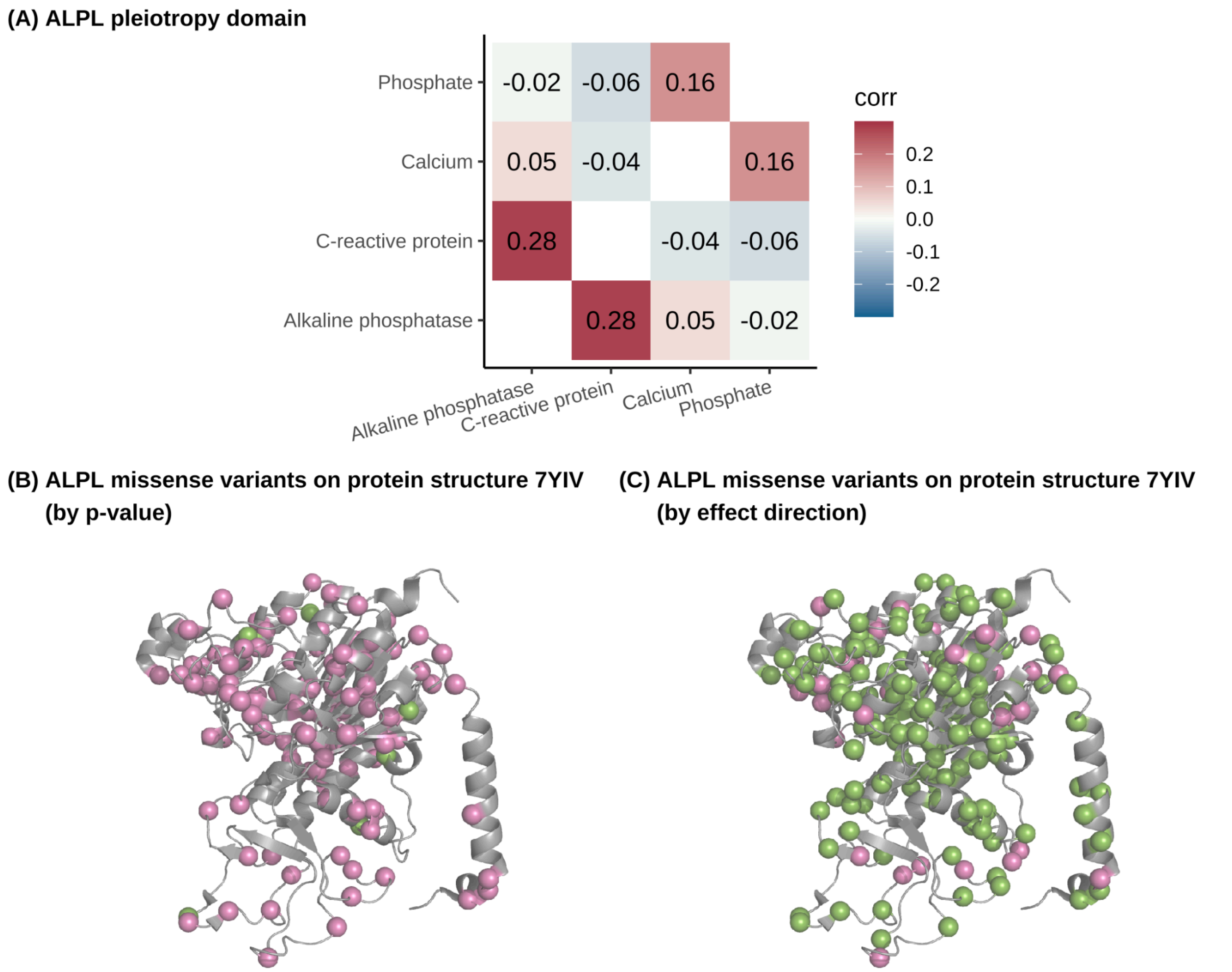


**Figure S17** | Cross-phenotype associations with *ALPL*. **A,** Correlation matrix of four phenotypes associated with *ALPL* across functional groups*.* **B,** Distribution of *ALPL* missense variants associated with alkaline phosphatase levels (pink) or phosphate levels but not alkaline phosphatase levels (green), mapped onto the 3D structure of ALPL protein (PDB ID: 7YIV). **C,** Distribution of *ALPL* missense variants affecting alkaline phosphatase and phosphate levels in the same direction (pink) or opposite directions (green), mapped onto the 3D structure of ALPL protein (PDB ID: 7YIV).

To assess the sensitivity of results to the allele frequency threshold, we applied ALLSPICE to the same set of association pairs across ultra-rare variants (AC < 5). The overall distribution of ALLSPICE test results is similar between rare (**Figure S15**; **SuppTable 5**) and ultra-rare (**Figure S18**; **SuppTable 6**) variants, although analyses restricted to ultra-rare variants show a better-controlled false positive rate at the cost of reduced statistical power. Among these

11,791 pairs of associations, 604 showed nominal evidence of heterogeneous variant effects ($p_{ALLSPICE} < 0.05$), including 340 pLoF, 216 missense, and 48 synonymous pairs. After correction for multiple testing ($p_{ALLSPICE} < 4.24 \times 10^{-6}$; 0.05/11,791 tests), 66 pairs remained significant (41 pLoF, and 25 missense; **Figure S18**). Consistent with observations for rare *ALB* variants (AF < $1 \times 10^{-4}$), ultra-rare *ALB* missense variants associated with calcium level are significantly closer to the calcium-binding sites on the ALB 3D protein structure than variants associated exclusively with albumin level ($P_{protein} = 3.42 \times 10^{-4}$; **Figure S19A** and **B**). In contrast, unlike rare variants, ultra-rare missense variants affecting albumin- and calcium-levels in opposite directions do not show a significant difference in distance to the calcium-binding sites ($P_{protein} = 0.054$; **Figure S19C** and **D**).

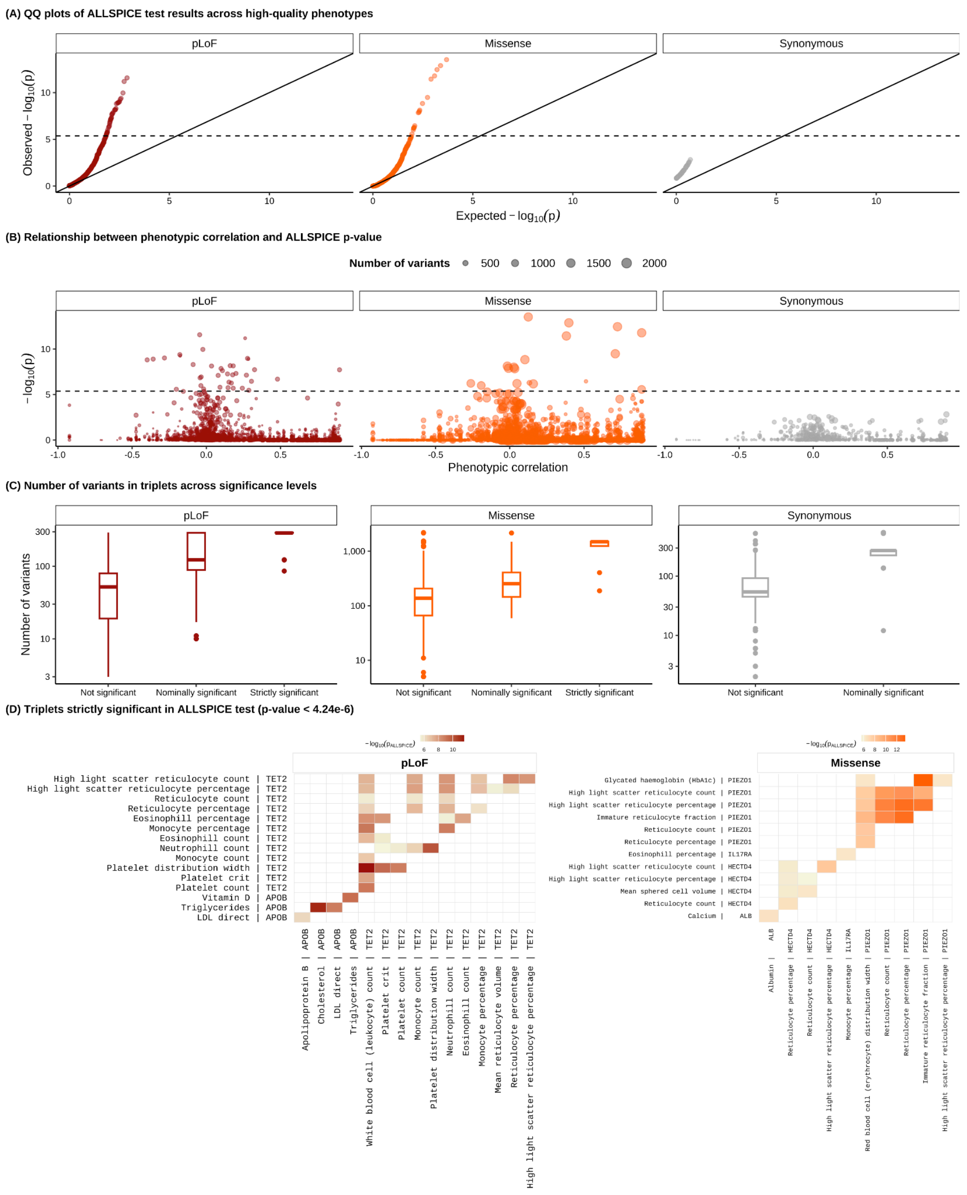


**Figure S18** | Summary of 11,791 ALLSPICE results across 359 continuous phenotypes on ultra-rare variants with AC < 5, restricted to association pairs with phenotypes having phenotypic correlation r < 0.9. **A,** QQplot of $p_{ALLSPICE}$ stratified by functional annotation group (colors and columns). The horizontal dashed line denotes the Bonferroni-corrected significance threshold (p-value=4.24 $\times 10^{-6}$). **B,** Relationship between phenotypic correlation (x-axis) and $p_{ALLSPICE}$ (-$\log_{10}$p, y-axis). Each point represents a single pair; columns indicate functional annotation group, and point size reflects the number of variants included in each test. **C**, Boxplots showing number of variants ($\log_{10}$-scaled y-axis) across three significance categories (x-axis): not significant ($p_{ALLSPICE} \geq 0.05$), nominally significant ($4.24 \times 10^{-6} \leq p_{ALLSPICE} < 0.05$), or strictly significant ($p_{ALLSPICE} < 4.24 \times 10^{-6}$), stratified by functional annotation group (colors and columns). **D**, Heatmap of $p_{ALLSPICE}$ for strictly significant association pairs: with phenotype pairs on the x- and y-axes (genes labeled accordingly). Cell color indicates -$\log_{10}$($p_{ALLSPICE}$), shown separately by functional annotation group.

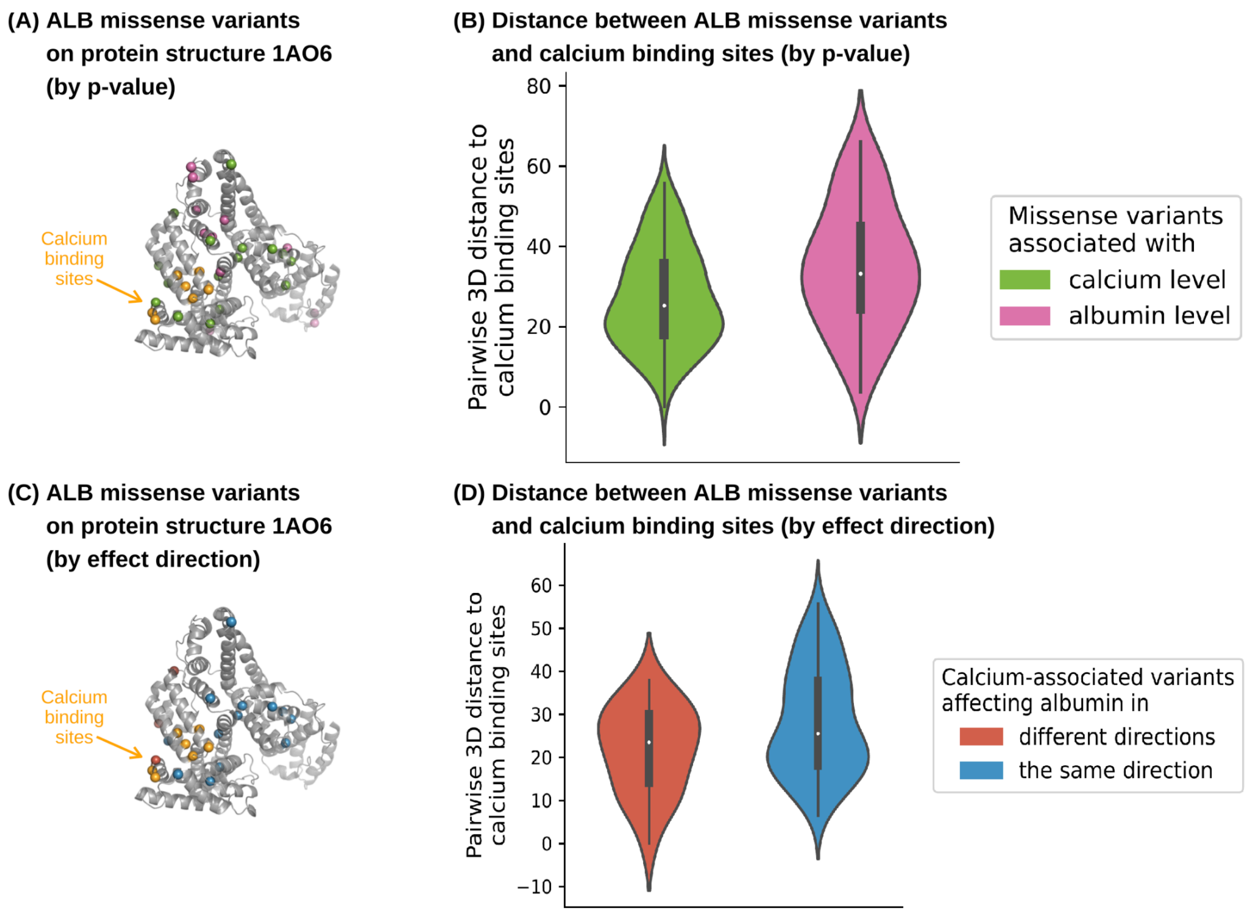


**Figure S19** | Ultra-rare *ALB* missense variants (AC < 5) on ALB protein structure. **A,** Distribution of *ALB* missense variants associated with calcium level but not albumin level (pink), associated with albumin but not calcium level (green), and calcium binding sites (orange) mapped onto the 3D structure of ALB protein (PDB ID: 1AO6). **B,** Comparison of pairwise distances from missense variants associated exclusively with calcium- (pink) or albumin- (green) levels to calcium binding sites on ALB protein structure in the 3D space. Colors in panel **B** are defined the same as in panel **A. C,** Distribution of *ALB* missense variants associated with either albumin or calcium level that affect the two biomarker levels in the same (blue) or opposite (red) direction(s), together with calcium binding sites (orange) on the 3D structure of ALB protein (PDB ID: 1AO6). **D,** Comparison of pairwise distances from missense variants to calcium binding sites on the ALB protein structure in the 3D space. Colors in panel **D** are defined the same as in panel **C.**

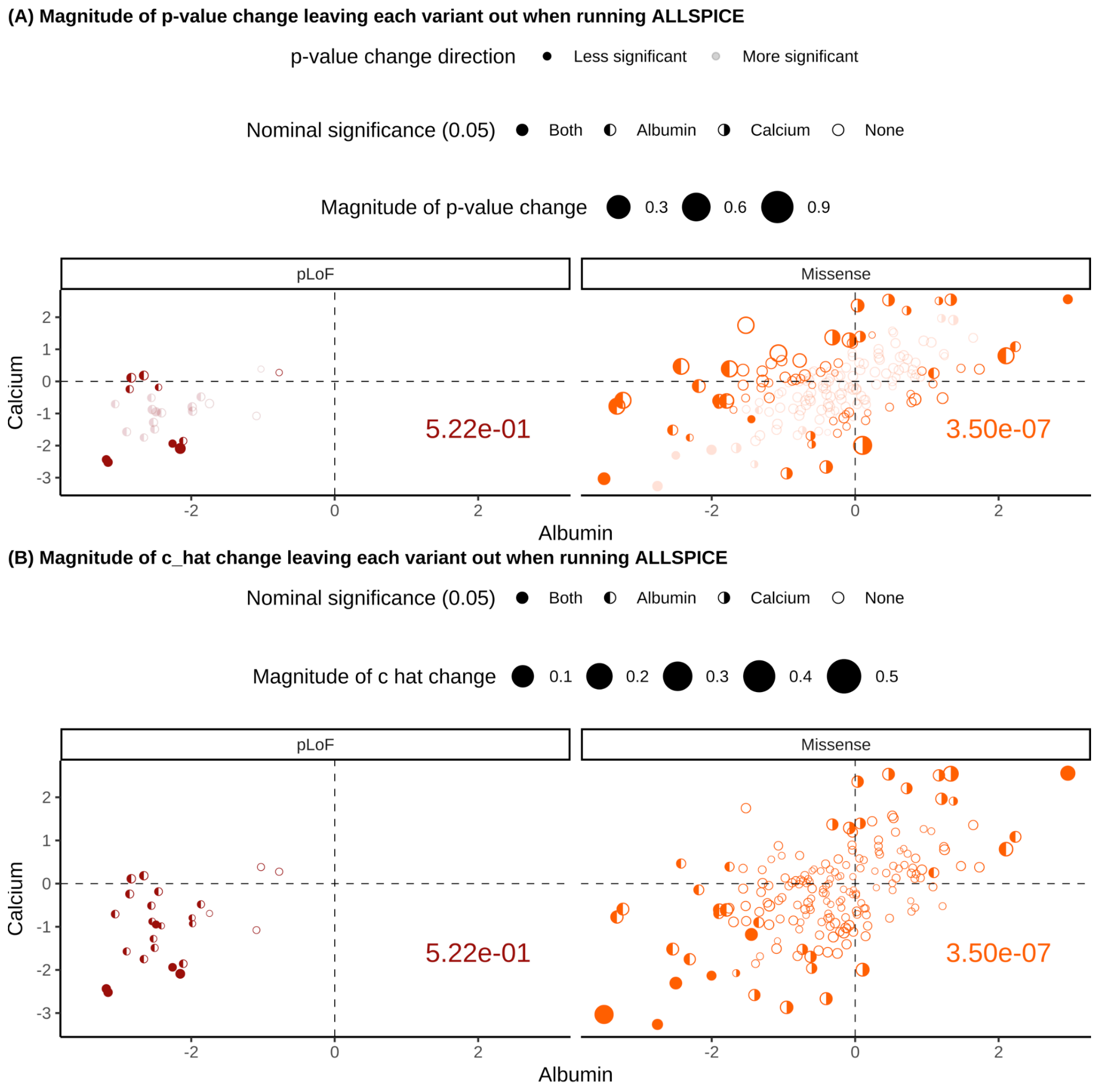


**Figure S20** | Leave-one-variant-out (LOVO) cross-validation results for ultra-rare variants in *ALB* (AC < 5). Each panel shows the comparison of variant effect sizes on albumin and calcium levels within *ALB*, stratified functional annotation group (colors and columns). Each point represents a variant in *ALB*. The x- and y-axes show the effect sizes of variants for albumin and calcium levels, respectively. Point shapes indicate the nominal significance of single-variant association tests ($p_{GWAS}$ < 0.05). The numbers in the bottom right of each panel report the $p_{ALLSPICE}$ for the corresponding annotation group, computed using rare variants with AC < 5. Point size reflects the influence of each variant in the LOVO analysis: **A** $|log_{10}(c_{LOVO}/c_{ALLvariants})|$ and **B** $|log_{10}(P_{LOVO}/P_{ALLvariants})|$ **.** In panel **A**, transparent points indicate variants whose removal increases test significance, whereas solidpoints indicate variants whose removal decreases significance.

References

1. Karczewski, K. J. *et al.* Systematic single-variant and gene-based association testing of thousands of phenotypes in 394,841 UK Biobank exomes. *Cell Genom* **2**, 100168 (2022).
2. Genovese, G. *et al.* Clonal hematopoiesis and blood-cancer risk inferred from blood DNA sequence. *N. Engl. J. Med.* **371**, 2477–2487 (2014).
3. Karczewski, K. J. *et al.* The mutational constraint spectrum quantified from variation in 141,456 humans. *Nature* **581**, 434–443 (2020).
4. Lek, M. *et al.* Analysis of protein-coding genetic variation in 60,706 humans. *Nature* **536**, 285–291 (2016).
5. Zeng, J. *et al.* Signatures of negative selection in the genetic architecture of human complex traits. *Nature Genetics* **50**, 746–753 (2018).
6. Gazal, S. *et al.* Functional architecture of low-frequency variants highlights strength of negative selection across coding and non-coding annotations. *Nature Genetics* **50**, 1600–1607 (2018).
7. O'Connor, L. J. *et al.* Extreme polygenicity of complex traits is explained by negative selection. *Am. J. Hum. Genet.* **105**, 456–476 (2019).
8. Szklarczyk, D. *et al.* STRING v11: protein-protein association networks with increased coverage, supporting functional discovery in genome-wide experimental datasets. *Nucleic Acids Res.* **47**, D607–D613 (2019).
9. von Mering, C. *et al.* STRING: known and predicted protein-protein associations, integrated and transferred across organisms. *Nucleic Acids Res.* **33**, D433–7 (2005).
10. Watanabe, K. *et al.* A global overview of pleiotropy and genetic architecture in complex traits. *Nat. Genet.* **51**, 1339–1348 (2019).
11. Levin, M. G. *et al.* Genome-Wide Assessment of Pleiotropy Across >1000 Traits from Global Biobanks. *medRxiv* (2025) doi:10.1101/2025.04.18.25326074.
